# Macrophage differentiation requires DNA damage caused by Caspase-Activated DNase

**DOI:** 10.1101/2023.10.16.562503

**Authors:** Deepak Maurya, Gayatri Rai, Debleena Mandal, Bama Charan Mondal

## Abstract

Phagocytic macrophages are crucial for innate immunity and tissue homeostasis. Most macrophages develop from embryonic precursors that populate every organ before birth to self-renew lifelong. However, the mechanisms for macrophage differentiation remain unknown. Using in vivo genetic analysis of the *Drosophila* larval hematopoietic organ, the lymph gland, we show that the developmentally regulated transient activation of Caspase-Activated DNase (CAD)-mediated DNA breaks in intermediate progenitors is essential for macrophage differentiation. Insulin receptor-mediated PI3K/Akt signaling triggers apoptotic signaling and causes DNA breaks during macrophage development. However, the same Akt signaling attenuates Apoptosis signal-regulating kinase 1 (Ask1) to control apoptotic and JNK activity in differentiating macrophages. DNA-damaged differentiating cells display autophagy activity as a survival strategy. Furthermore, caspase activity is required for embryonic-origin macrophage development and efficient phagocytosis. This study reveals a previously unknown relationship between developmental signals and caspase-activated DNA breaks necessary for multifunctional macrophage differentiation.

## Introduction

Macrophages are evolutionarily conserved phagocytic cells with crucial roles in innate immunity, development, tissue-specific function, and monitoring of aberrant cells like those of cancer^1, 2^. A comprehensive understanding of macrophage development is essential, as manipulating macrophages for clinical use is gaining popularity^3^.

Fate mapping, single-cell transcriptomics, and epigenetic studies showed that heterogeneous tissue-resident macrophages arise from early embryonic (yolk sac and fetal liver) erythro-myeloid progenitors and reside lifelong with limited self renewal^4–7^. However, in some tissues like the intestine, the bone marrow-derived circulating monocytes differentiate into tissue-specific macrophages when needed^8^. Myeloid progenitor differentiation requires precise control of gene expression, which is regulated by transcription factors, chromatin landscape, cellular metabolism, autophagy, apoptotic factors, and systemic cues during development and disease^9–14^. Apoptotic signaling activates protease caspases that target hundreds of proteins during cell death^15^. Studies have also shown that caspases are activated during various types of cell differentiation^16–18^. Interestingly, activated caspases are required to develop mammalian myeloid cells, such as erythrocytes, platelet development, and CSF1-mediated monocyte-to-macrophage differentiation^13^. However, the precise mechanism responsible for macrophage differentiation remains unknown.

To investigate the mechanisms underlying macrophage heterogeneity and versatility, we use in vivo genetic analysis of the *Drosophila* hematopoietic system, which has myeloid-type blood cells only. This system uses evolutionarily conserved molecular factors in development and innate immunity^19, 20^. Hematopoiesis in *Drosophila* occurs in two waves. First-wave blood cells (hemocytes) develop in the early embryo’s head mesoderm, contributing to embryonic, larval, and adult stages. The second wave begins at the late embryonic stage in the cardiogenic region to generate the larval hematopoietic organ called the lymph gland, which resembles a mammalian hematopoietic organ with a niche, multipotent progenitors, intermediate progenitors or differentiating cells, and differentiated cells (**Figure 1A**)^19–21^. Blood progenitors are housed in the inner core of the lymph gland and proliferate during the early larval stages. At the mid-second instar, they stop dividing and begin to differentiate into plasmatocytes and crystal cells at the distal margins in the lymph gland’s outer boundary, which disperse during pupation and contribute to the adult blood cells. Most blood cells in *Drosophila* are macrophage-like cells called plasmatocytes (hereafter referred to as macrophages) that function similarly to mammalian macrophages^22, 23^. Recent studies using enhancer analysis^24^ and single-cell transcriptomics suggest that *Drosophila* larvae and adults have heterogeneous tissue-specific macrophages like vertebrates^22, 25^.

**Figure 1.**
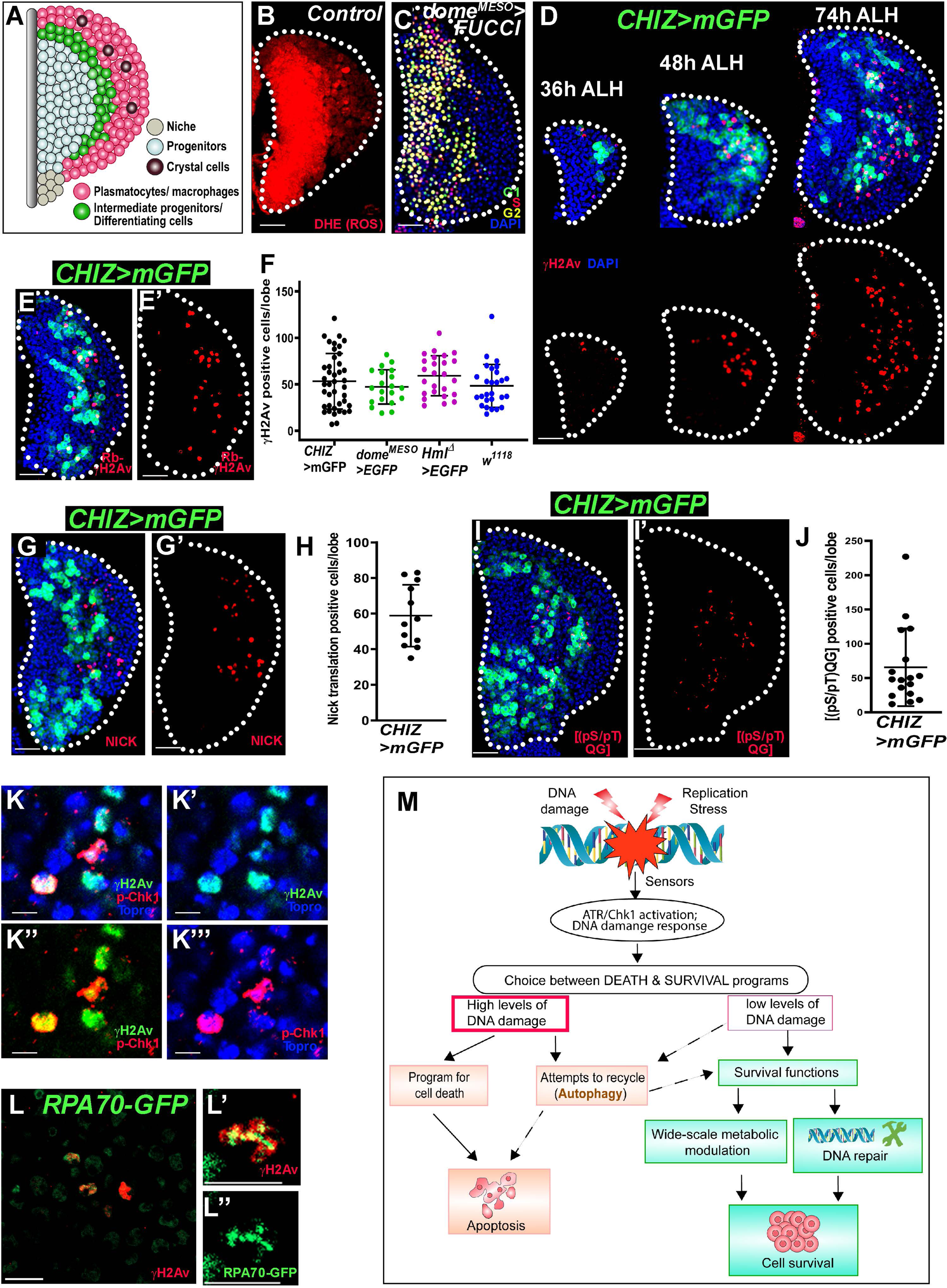
DNA damage occurs during the differentiation of lymph gland. **(A)** Schematic of the third instar primary lymph gland lobe showing locations of different cell types. **(B)** Dihydroethidium (DHE) staining (red) in control *dome^MESO^-Gal4 UAS-2xEGFP/+* (GFP channel not shown) displays high Reactive Oxygen Species (ROS) in lymph gland progenitors. **(C)** High number of G2 cell cycle phase (yellow) progenitors in *dome^MESO^-Gal4*, *UAS-FUCCI*. **(D)** DNA damage response (DDR) marker immunostaining of mouse anti-γH2Av (red) in control lymph glands (*CHIZ-Gal4 UAS-mGFP/+* background) at 38h After Larval Hatching (ALH), 48h ALH, and 74h ALH show increased number of γH2Av positive cells with larval age in the intermediate zone marked CHIZ>mGFP (green) — a high magnification γH2Av staining shown in Figure S1. **(E-E’)** Staining of a rabbit anti-γH2Av (red) antibody in *CHIZ-Gal4 UAS-mGFP*/*+* background with CHIZ+ (green) (E) and only γH2Av (E’) show a similar staining pattern with mouse anti-γH2Av antibody (D, 74h ALH). **(F)** Quantification of γH2Av positive cell number in different genetic backgrounds: *CHIZ-Gal4 UAS-mGFP/+* (n=41), *dome^MESO^-Gal4 UAS-2xEGFP/+* (n=20), *Hml^Δ^-Gal4 UAS-2xEGFP /+* (n=25) and *w^1118^* (n=27) per lymph gland lobe and images are shown in Figure S1. **(G-G’)** Nick translation (red) positive cells are seen in the intermediate zone of control lymph gland *CHIZ-Gal4 UAS-mGFP/+* (n=12), indicating DNA strand breaks. **(H)** Quantification of DNA nick translation positive cells of lymph gland represented in (G-G’). **(I-I’)** Immunostaining of phospho-ATM/ATR substrate motif [(pS/Tq) QG] (red) show in intermediate zone (*CHIZ>mGFP/+*) (n=17), indicating ATM/ATR activity as DDR in control lymph gland. **(J)** Quantification of ATM/ATR substrate motif positive cells of lymph glands represented in (I-I’). **(K-K’’’)** High magnification image of the control lymph gland shown with a well-known DDR marker p-Chk1 (red) colocalizes in the same cells with γH2Av (green), and nuclei mark Topro3 (blue) (K), γH2Av and Topro3 (K’), γH2Av and p-Chk1 (K’’), p-Chk1 and Topro3 (K’’’). **(L-L’’)** DDR protein RPA1 homolog RPA70; high intensity of *RPA70-GFP* (green) colocalize with γH2Av (red) positive cells (whole lymph gland lobe shown in Figure S1G-S1G’). **(M)** A schematic showing the choice between cell death (apoptosis) or the survival process upon DNA damage. All images are from wandering third instar lymph gland except image D, where the lymph glands are at 38h ALH, 48h ALH, and 74h ALH stages. Scale bars: 25µm in all images except 10µm in (L) and 5µm in (L’-L’’) with maximum intensity projections of the middle third optical sections except image (B, C, K-L’’), which are single optical sections. Nuclei stained with DAPI (blue), except (K). Lymph glands boundary demarcated by white dotted line for clarity. Error bars, mean ± standard deviation (SD). All images are representative of 3 or more independent biological experiments.

The lymph gland undergoes a developmental program regulated by local signals from its microenvironment^26^, cell-autonomous factors downstream of Pvr, and JAK/STAT signaling^27–, 29^, or systemic signals (e.g., InR, GABA-R)^30–32^. Recent report indicates that maintenance of progenitors in lymph glands by Wnt6-EGFR-mediated signaling^33^. Besides, the third-instar lymph gland progenitors show a high level of Reactive Oxygen Species (ROS)^34^ (**Figure 1B**), akin to mammalian common myeloid progenitors, but have a lengthy G2 cell cycle phase (**Figure 1C**). Interestingly, G2 arrest occurs during stress-mediated DNA breakage that triggers the DNA damage response (DDR), leading to arrest until the damage is adequately repaired or apoptosis occurs^35^.

Here, we show that a sublethal apoptotic caspases activate Caspase-Activated DNase (CAD), triggering DNA breaks during the normal development of *Drosophila* macrophages. We find that insulin receptor-mediated PI3K/Akt signaling in differentiating macrophages induce sublethal caspase activation by attenuating Ask1. Furthermore, caspase activity is required during embryonic-origin and lymph gland macrophages development for efficient phagocytosis. This study provides insight into developmental signaling and caspase-activated DNA breaks in differentiating macrophages that promote macrophage-specific gene expression required for adaption to tissue-specific microenvironments and for efficient innate immune responses.

## Results

### DNA breaks occur during myeloid-type progenitor cell differentiation in lymph glands

We investigated whether developmentally controlled G2 arrest in the third instar lymph gland progenitors (**Figure 1C**) is due to DNA damage. We first monitored the status of DDR marker-positive cells using a mouse anti-γH2Av (γH2AX homolog) antibody^36^ during larval development in the lymph gland cells^37, 38^. Interestingly, we found that γH2Av-positive cells appear in the periphery of the mid-second instar [36 hours after larval hatching (ALH)] lymph gland, which coincides with the onset of differentiation^28^ (**Figure 1D**). The γH2Av positive cell numbers increase in the differentiating zone during the early third instar (48 hours ALH). By the wandering third instar stage (74 hours ALH), the number of the differentiating zone located γH2Av positive cells further increased (**Figure 1D**).

Differentiating cells or intermediate progenitors co-express the progenitor marker d*ome^MESO^* and the earliest differentiating cell marker *Hml*^39^. Using the *split-Gal4* strategy, a driver *CHIZ-Gal4 UAS-mCD8::GFP* (hereafter *CHIZ>mGFP* genotype or CHIZ^+^ cells) was made^39^ (green cells in **Figures 1A**, **1D**) that could mark most of the differentiating or intermediate progenitor zone. Similar to Figure 1D, most of the γH2Av positive cells in the differentiating zone (CHIZ^+^) (**Figures 1E-1E’**) were also found with another widely used rabbit anti-γH2Av antibody^40^. Notably, γH2Av staining covered the entire nuclear region, except the DAPI-bright heterochromatin region (**Figures S1A-S1A”’**).

A connection between differentiating cells and DDR is an important finding and was therefore confirmed in multiple genotypes using several different strategies. We assessed the status of γH2Av-positive cell numbers in fly lines used to study *Drosophila* hematopoiesis, such as *w*^*1118*^, *CHIZ>mGFP, Hml^Δ^-Gal4, UAS-2xEGFP, and dome^MESO^-Gal4, UAS-2xEGFP*. The third instar lymph glands of all the examined genotypes showed a similar number of γH2Av positive cells in the differentiating zone (**Figures 1F** and **S1B-S1D**); thus excluding the possibility of a genetic background effect. We evaluated DNA breakage with an *in vivo* nick translation assay and found incorporation of DIG-labeled dTTP, which indicates DNA repair synthesis^41^. The number of DIG-labeled nuclei was similar to γH2Av-labeled nuclei in the intermediate zone (**Figures 1G-1G’, 1H** and **S1E-S1F**). Upon DNA damage, ATR/ATM kinases are activated by phosphorylation, which further phosphorylates H2Av, Chk1, and other proteins involved in DDR and DNA repair^35^. Immunostaining for the phospho-ATM/ATR substrate motif [(pS/pT)QG]^40^ revealed the lymph glands staining pattern to be similar to the γH2Av positive cells (**Figures 1I-1I’** and **1J**). Phospho-Chk1, a well-known DDR marker^42^, also co-localized with γH2Av-positive cells in the lymph gland (**Figures 1K-1K”’**). RPA1 homolog RPA70 is involved in DDR^43^. Using the *RPA70-GFP* line^44^ and immunostaining for γH2Av, we found that the high-intensity RPA70-GFP puncta co-localized with γH2Av in the lymph gland (**Figures 1L-1L”** and **S1G-S1G’**). Together, these confirm that DNA damage repair foci are widely present in a subset of intermediate progenitors in the lymph gland.

We employed the Fluorescence Ubiquitin Cell Cycle Indicator (FUCCI) system^45^ to assess the cell cycle status of lymph gland cells that show DNA damage. We used the *e33c-Gal4* driver to identify G1, S, G2, and M phases in the entire lymph gland in the third instar larvae and found that all the γH2Av-positive cells were in the G2 phase (**Figures S1H-S1H””**). These DDR-marked cells were different from the PCNA-GFP^46^ positive cells, an S phase marker (**Figures S1I-S1I’**). However, the nuclear pore complexes marked by Nup98-GFP remained intact in γH2Av positive cells (**Figures S1J-J”’**).

These findings thus indicate that developmental DNA damage in the intermediate progenitors is a feature of the differentiating myeloid-type blood cells. Therefore, the next question addressed in this study was to identify the developmental cues involved in causing DNA damage during myeloid-type cell differentiation.

### Caspases activation and DNA breaks in differentiating myeloid-type blood cells

Cells with damaged nuclear DNA activate various factors, including damage sensors, DDR, cell cycle checkpoints, and DNA repair proteins for cell survival. The cell either adopts cell death (apoptosis) fate or activates the survival process depending on the strength of damage signals (schematic in **Figure 1M**)^35^. We first tested whether apoptosis pathways are activated in the third instar lymph gland. To check the *Drosophila* executioner caspase Dcp-1 activity (schematic in **Figure 2A**)^17^, we used the cleaved *Drosophila* Dcp1 (Asp215) antibody^47^ (CST, USA, Cat#9578S) immunostaining of third instar lymph gland. Remarkably, we detected cleaved Dcp-1 (hereafter, Dcp1) positive cells in the intermediate zone (**Figures 2B-2B’,** and quantification showed in **2C**) in *CHIZ>mGFP* heterozygous genetic background. Significantly, more than 90% of γH2Av-positive cells were also Dcp-1-positive cells (**Figures 2D-2D”** and **2E**). However, the number of Dcp-1 positive cells was higher than the γH2Av positive cells (**Figure S2A**), indicating that caspases activate first to initiate DNA breakage during lymph gland development. Our finding that the intermediate zones in *dome^MESO^-Gal4, UAS-2xEGFP,* and *w*^*1118*^ genotypes also have similar frequencies of caspase-positive cells (**Figures 2C, S2B-S2C, S2D**) rule out the possibility of a genetic background effect.

**Figure 2.**
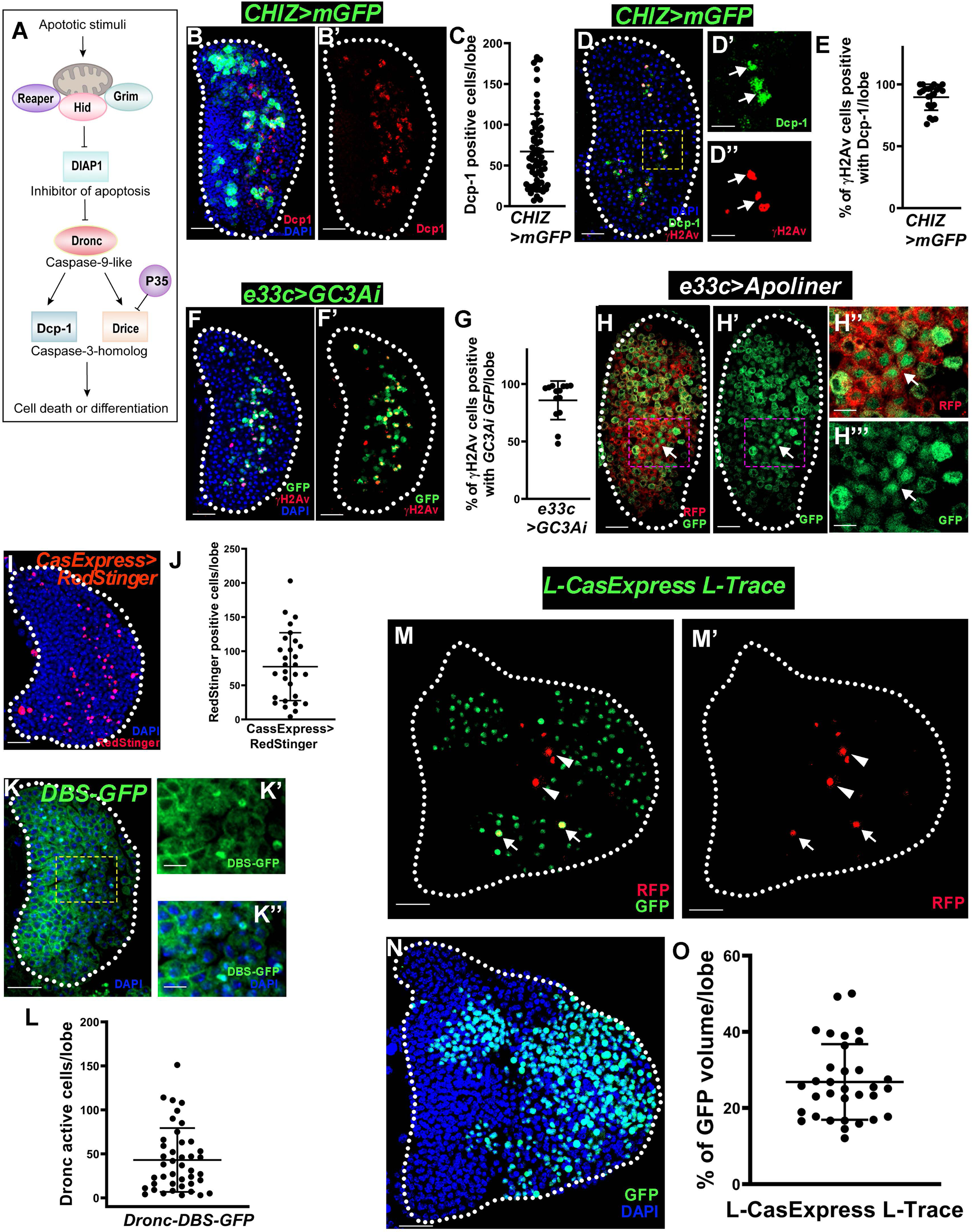
DNA breaks and active caspase in differentiating progenitor blood cells. **(A)** A schematic model showing the *Drosophila* apoptotic pathway. **(B-B’)** Intermediate progenitor zone of control [*CHIZ>mGFP/+* (green)] lymph gland exhibits immunostaining of cleaved Dcp-1 (red). **(C)** Quantification of Dcp-1 positive cell numbers in *CHIZ>mGFP/+* (n=66) genetic background per lymph gland lobe. **(D-D’’)** γH2Av (red) cells are also Dcp-1 (green) positive in *CHIZ>mGFP/+* lymph gland showing without CHIZ>mGFP (D), inset show Dcp-1 (C’) and γH2Av (C’’). **(E)** Quantification shows >90% percentage of γH2Av positive cells colocalized with Dcp-1 positive cells in control lymph glands per lobe [*CHIZ>mGFP/+* (n=22)]. **(F-F’)** GFP (green) fluorescent reporter of executioner caspase activity (*e33c-Gal4, UAS-GC3Ai*) shows caspase activity which colocalizes with γH2Av staining (red) (n=14). **(G)** Quantification of (F-F’) showed the percentage of colocalization per lymph gland lobe. **(H-H’’’)** Intermediate zone of lymph gland shows caspase activity by expression of Apoliner (*e33c-Gal4*, *UAS*-*Apoliner*) a membrane RFP (red) and nuclear GFP (marked caspase active cell) (H), only GFP (H’). Insets show high magnification of membrane RFP and membrane as well as nuclear GFP (H’’) and only GFP (H’’’). The arrow indicates caspase active cells. **(I)** Expression of *CasExpress-Gal4*, *UAS-RedStinger* (n=29) shows executioner caspase activity (red) in the lymph gland. **(J)** Quantification of caspase active cells per lymph gland lobe represented in (I). **(K-K’’)** Initiator caspase Dronc activity shown by nuclear *Drice-Based-Sensor-GFP (DBS-GFP)* (n=42) in the lymph gland intermediate zone (K). High magnification images showing nuclear GFP (green). Colocalization with γH2Av is shown in Figure S2I. **(L)** Quantification of *DBS-GFP* cells per lymph gland lobe represented in (K). **(M-N)** The *L-CasExpress L-Trace* (*lex-Aop-Flp::Ubi-FRT-STOP-FRT-GFP/lex-Aop-2XmRFP; L-caspase/+*) shows real-time executioner caspase activity in RFP (red) cells marked by the arrowhead (M-M’), with caspase lineage trace cells with GFP (M-M’), colocalized cells marked by the arrow, middle 3^rd^ section lineage trace GFP (green) (N). **(O)** Quantification of the ratio of caspase lineage cells and DAPI volumes in the *L-CasExpress L-Trace* (n=33) per lymph gland lobe. All images are single optical sections except images (B-B’, I, and N) which are maximum intensity projections of the middle third optical section of the wandering 3^rd^ instar lymph gland. Scale bars: 25µm in all images except 5µm in (H’’-H’’’) and 10 µm in (K’-K’’). Nuclei stained with DAPI (blue). Lymph glands boundary demarcated by white dotted line for clarity. Error bars, mean ± SD. All images are representative of 3 or more independent biological experiments.

We confirmed the executioner caspase activity in the lymph gland through multiple approaches. First, using *UAS-GC3Ai,* a fluorescent sensor of executioner caspase activity^48^, we found high caspase activity only in the differentiating zone (**Figures 2F-2F’** and for quantification **2G**). Likewise, *e33c-GAL4* driven^26^ *UAS-VC3Ai* activity (**Figures S2E-S2E””**) and γH2Av positive cells co-localized with caspase active cells (**Figure 2F-2F’**). We also used the Apoliner caspase reporter, *UAS-Apoliner* expressed using both *e33c-Gal4* and *CHIZ-Gal4* drivers where mRFP and GFP are initially membrane-bound, but upon caspase activation GFP translocates to the nucleus^49^. We observed nuclear GFP in the differentiating region (**Figures 2H-2H’’’**) and in a subset of CHIZ+ cells (**Figures S2F-S2F’**). Further, we used *CasExpress-Gal4 (*BL65420)^47^ and *UAS-RedStinger* (BL8546) reporters that showed executioner caspase-positive cells in the intermediate zone (**Figure 2I** and for quantification **2J**) but not in the mutant form of *CasExpress^mutant^-Gal4 (*BL65419)^47^ (**Figure S2G**). Published literature suggests that high levels of TUNEL-positive cells have no return from cell death^50^. Thus, to test whether γH2Av-positive cells in the lymph gland are high-intensity TUNEL-positive cells, we performed TUNEL staining. Remarkably, γH2Av-positive cells in the lymph gland are not high-intensity TUNEL-positive cells (**Figures S2H-S2H”**), though some high-intensity TUNEL-positive cells present in the differentiated zone. This result indicates that γH2Av-positive cells are not dying. Collectively, these results suggest that executioner caspase activity is sublethal in differentiating cells in the lymph gland.

In *Drosophila*, activated initiator caspase Dronc cleaves executioner caspases Drice and Dcp-1. Dronc is also known to have non-apoptotic functions in various tissues^51^. Therefore, we checked Dronc activity using a Drice-based sensor (DBS) line^52^. Interestingly, third-instar lymph glands show nuclear-localized Histone-GFP (DBS) in the intermediate zone (**Figures 2K-2K”** and **2L**), which co-localizes with γH2Av staining that suggests Dronc is also active in this zone. However, we observed that γH2Av-positive cells have lower DBS intensity than only DBS cells (**Figures S2I-S2I”**). Thus, this result hints that initiator caspase Dronc activated along with DDR only in a subset of differentiating cells because of their temporal activity.

Finally, we traced the lineage of executioner caspase-activated cells in the lymph gland using a new caspase lineage trace marker line L-CasExpress L-Trace^53^. Briefly, a membrane-bound LexA is cleaved upon executioner caspase activation and transported to the nucleus to bind lexAOP regulators. The caspase active cells could be marked in real-time by nuclear RFP, while the flippase expression caused somatic recombination in the same cells so that nuclear RFP positive cells were permanently marked progeny cells with nuclear GFP expression (a schematic showed in **Figure S2J**)^53^. Remarkably, the third instar lymph gland showed RFP-positive cells in the intermediate zone, while the lineage trace GFP-positive cells were distributed throughout the entire differentiated zone (**Figures 2M-2N**). Quantification of caspase lineage cells (GFP+) showed them to be 27% of total lymph gland cells (**Figures 2N-2O**), a value similar to the number of hemolectin-positive cells present in the lymph gland differentiated zone shown by Spratford et al. 2021^39^. Only 9% of the crystal cells are caspase lineage trace positive (**Figures S2K-S2K’** and quantification in **S2L**). Of note, crystal cells could differentiate from immature macrophages depending on active Notch singling^54^. Thus, this indicates that the differentiation of crystal cells is independent of caspase activation. These results show that differentiating cells with transiently activated executioner caspases in the lymph gland survive so that the differentiated zone becomes full of macrophages.

### Caspase-mediated DNA damage is required for macrophage differentiation

We next checked the involvement of the *Drosophila* apoptotic pathway (**Figure 2O**) in macrophage differentiation. Firstly, we examined numbers of γH2Av-positive cells in mutants of *Drosophila* executioner caspase (*Drice*^*2c8*^*/Drice^Δ1^*)^55, 56^ and initiator caspase (*Dronc^I^*^24^*/Dronc^I^*^29^)^57^, and found severely low numbers of γH2Av-positive cells (**Figures 3A-3C’,** quantitation showed in **3D**), and macrophage marked by the phagocytic receptor Draper^58^ (**Figures 3E-3G’,** quantification showed in **3H**)^59^. In *Drice* and *Dronc* mutants, we observed that the active Dcp-1 positive cells were absent (**Figures S3H-S3J’**, quantification in **S3K**). Together, these suggest that caspases regulate lymph gland progenitor differentiation.

**Figure 3.**
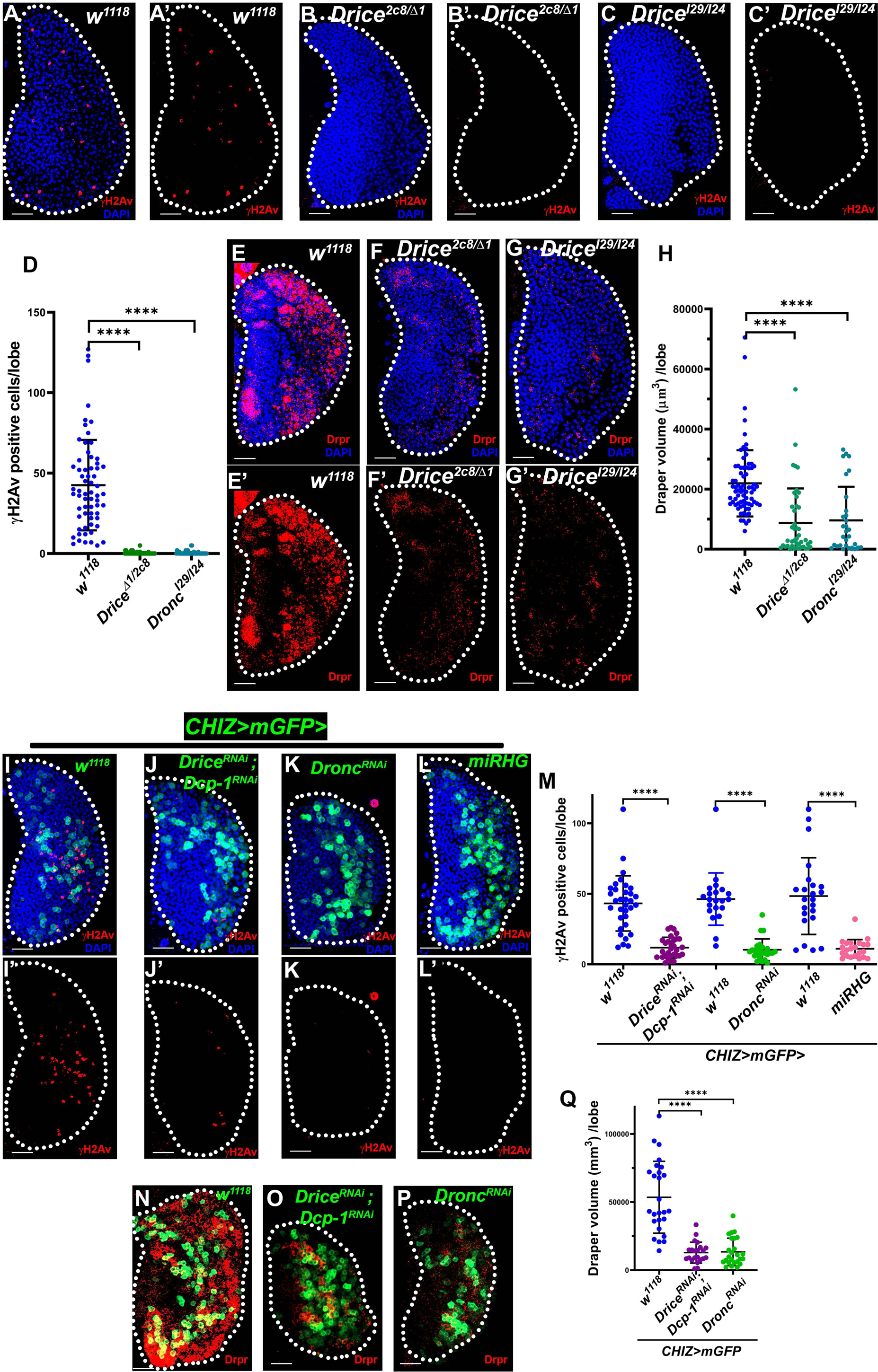
Caspase-mediated DNA breaks are necessary for macrophage differentiation. **(A-C’)** γH2Av staining (red) is shown in control *w^1118^* (n=74) (A-A’) lymph gland, but in the mutants of executioner *Drice^Δ1^/Drice^2c8^* (n=43) (B-B’) and initiator caspase, *Dronc^I29^/Dronc^I24^*(n=30) (CC’) do not show γH2Av positive cells. **(D)** Quantification of γH2Av positive cells of lymph gland represented in (A-C’). **(E-G’)** Phagocytic receptor Draper staining (red) as macrophage marker in control *w^1118^*(n=64) (E-E’), and mutants *Drice^Δ1^/Drice^2c8^* (n=71) (F-F’), *Dronc^I29^/Dronc^I24^* (n=51) (G-G’) shows severely less Draper staining. **(H)** Quantification of Draper volume of lymph gland represented in (E-G’). **(I-L’)** One representative image of control, *CHIZ>mGFP/+* (green) (I-I’), lymph gland lobe shown with *CHIZ>mGFP* driven *RNAi*, UAS-*Drice^RNAi^; UAS-Dcp-1^RNAi^* (J-J’)*, UAS-Dronc^RNAi^* (K-K’), and *UAS-miRHG* (L-L’) show less γH2Av positive cells (red) compared with control (I). **(M)** Quantification of γH2Av positive cells in control, *CHIZ>mGFP/+* (n=33) with *UAS-Drice^RNAi^; UAS-Dcp-1^RNAi^* (n=30); control, *CHIZ>mGFP/+* (n=21) with *UAS-Dronc^RNAi^* (n=24); and control, *CHIZ>mGFP/+* (n=23) with *UAS-miRHG* (n=22) show significant reduction in γH2Av positive cells. Here, control sets are different for each RNAi because one set of experiments with a particular RNAi line was done separately. **(N-P)** One representative image of control, *CHIZ>mGFP/+* (n=26) (N) with Draper staining (red), lymph gland lobe shown with *CHIZ>mGFP* driven *RNAi*, *UAS*-*Drice^RNAi^; Dcp-1^RNAi^* (n=22) (O) and *UAS-Dronc^RNAi^* (n=25) (P) show drastically lower Draper expression than the control. **(Q)** Quantification of Draper volume of lymph gland represented in (N-P). All images show a 25µm scale bar, maximum-intensity projections of the middle third optical section of the wandering third instar larval lymph gland lobe. Nuclei stained with DAPI (blue). Lymph glands boundary demarcated by white dotted line for clarity. ****P < 0.0001 Error bars, mean ± SD. All images are representative of 3 or more independent biological experiments.

To exclude the phenotypes in these caspase mutants larvae to be a consequence of systemic signals involved in lymph gland progenitor maintenance^30, 31^, we conducted further experiments to determine the apoptotic pathway role in the lymph glands. Blocking the apoptotic pathway in the intermediate progenitors by expressing the widely used microRNA against reaper, hid, and grim (RHG) transcripts (*UAS*-*miRHG*)^60^, resulted in significantly fewer γH2Av positive cells (**Figures 3I-3I’, 3L-3L’** and **3M**) and loss of caspase active cells (**Figures S3C-S3C’, S3F-S3F’** and **S3G**). Inhibiting executioner caspase by baculovirus protein P35^60^ caused a similar phenotype (**Figures S3A-S3B’**), although weaker than the miRHG phenotype. Further, depletion of both executioner caspases using RNA interference (RNAi) of *Drice* and *Dcp-1*^61^ in the intermediate progenitors resulted in significantly fewer DDR cells (**Figures 3I-3J**, quantification in **3M**), with substantially low phagocytic marker Draper positive macrophages (**Figures 3N-3O, 3Q**). Next, we knocked down Dronc in the intermediate progenitor using *Dronc^RNAi^* and found fewer γH2Av positive cells (**Figures 3I, 3K, 3M**) and significantly reduced Draper staining (**Figures 3N,3P-3Q**). Dcp1 caspase activity is reduced in both these caspase depletion backgrounds (**Figures S3C-S3E’**, quantification in **S3G**). These results clearly demonstrate that the *Drosophila* apoptotic signaling pathway, including initiator and executioner caspases, is required for the DNA breaks-mediated phagocytic macrophage differentiation.

### InR/PI3K/Akt signaling regulates caspase activity and DDR in macrophage differentiation

The crucial issue of the physiological relevance of the above results requires investigation of upstream mechanisms that trigger apoptotic signaling and potentially involve DNA damage during the normal development of macrophages. Studies showed that signaling level changes in the lymph gland can influence the behavior of blood progenitors^27, 28, 30, 62^. Thus, we knocked down candidate genes in the lymph gland by RNAi and monitored for the loss of caspase activity. This survey identified Akt as a signaling factor required for caspase activation in the lymph gland since Akt knockdown in the intermediate progenitors (*CHIZ>mGFP>Akt^RNAi^*) was associated with significantly fewer Dcp-1-positive cells (**Figures 4A-4B’, S4A-B’,** quantification showed in **4V**). We further investigated whether InR/PI3K/Akt-mediated signaling plays a role in this process. Reduction in the PI3K signaling in intermediate progenitors through expression of PI3K dominant negative form (*UAS-PI3K92E^DN^*) (**Figures 4A-4A’, 4C-4C’,** quantification showed in **4V**) and depletion of InR in the intermediate progenitors (*CHIZ>mGFP>InR^RNAi^*) resulted in significantly reduced number of Dcp-1 positive cells (**Figures 4A-4A’, 4D-4D’,** quantification showed in **4V**). This is consistent with a recent study that PI3K/Akt signaling activation induces autonomous apoptotic stress^53^. We also assessed γH2Av and Draper staining by expressing *InR^RNAi^*, A*kt^RNAi,^* and *PI3K^DN^* in intermediate progenitors and found that the number of γH2Av-positive cells dramatically decreased (**Figures 4H-4K’,** quantification showed in **4W**). In these backgrounds, Draper staining levels in lymph glands were significantly reduced (**Figures 4O-4R**, quantification showed in **4X**), similar to Draper expression in glia regulated by InR/Akt signaling^58, 63^. The lymph gland size was significantly reduced (**Figure S4C**) in these genetic backgrounds. However, the number of intermediate progenitors (CHIZ^+^) cells remained unchanged (**Figures S4D**), suggesting that blocking InR/PI3K/Akt signaling in intermediate progenitors causes the lymph gland differentiation halts. We also assessed crystal cells (involved in melanisation and blood clotting) numbers using Hindsight (Hnt)^19^ and differentiated macrophages numbers using P1 (also called NimC1)^19, 64^ as another marker. The number of crystal cells remained unaffected following depletion of Akt in the intermediate progenitors (**Figures S4E-S4F,** quantification **S4G**). However, the macrophage numbers decreased significantly (**Figures S4H, S4J,** quantification in **S4K**). Thus, these results show that inhibition of InR/PI3K/Akt signaling suppresses macrophage differentiation and caspase-mediated DNA damage.

**Figure 4.**
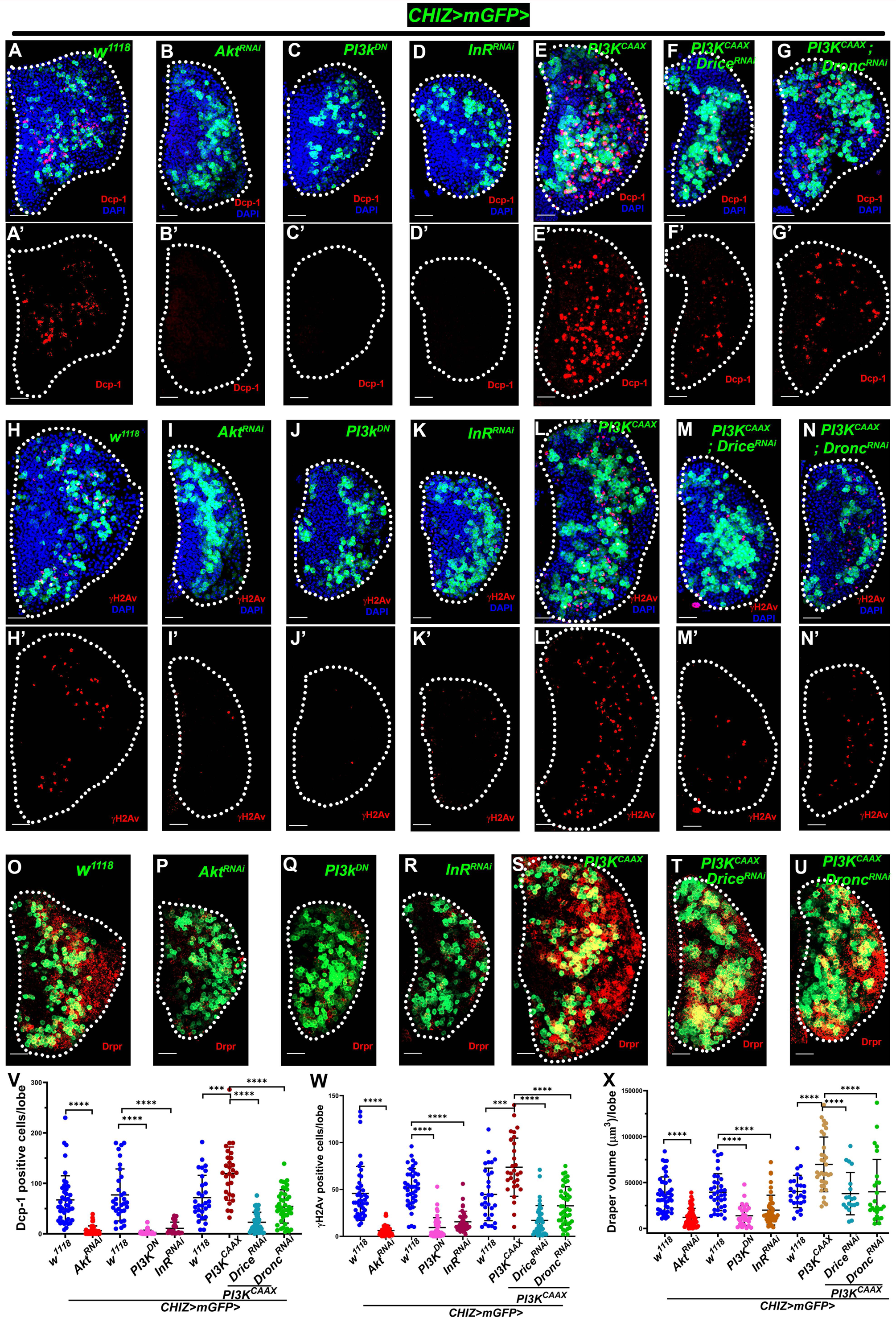
InR-PI3K-Akt signaling regulates caspase activity and DNA breaks in macrophage differentiation. **(A-G’)** InR/PI3K/Akt-mediated executioner caspase Dcp-1 regulation: Dcp-1 staining (red) in control, *CHIZ>mGFP/+* (n=105) (A-A’) (green), *CHIZ>mGFP* driven experimental sets [*Akt^RNAi^* (B-B’) (n=50), *PI3K^DN^* (C-C’) (n=37), *InR^RNAi^* (D-D’) (n=22)] show less Dcp-1 positive cells. Also, *CHIZ>mGFP* driven *PI3K^CAAX^* (E-E’) (n=29) have high Dcp-1 positive cells and are rescued in *PI3K^CAAX^; Drice^RNAi^* (F-F’) (n=42) and *PI3K^CAAX^*; *Dronc^RNAi^* (G-G’) (n=43). **(H-N’)** *CHIZ>mGFP* (green) driven experimental sets [*Akt^RNAi^* (I-I’) (n=47), *PI3K^DN^*(J-J’) (n=48), *InR^RNAi^* (K-K’) (n=36)] show less γH2Av positive cells (red) compared to control, *CHIZ>mGFP/+* (H-H’) (n=113); *CHIZ>mGFP* driven *PI3K^CAAX^* (L-L’) (n=29) show high γH2Av positive cells (red) and rescued in *PI3K^CAAX^; Drice^RNAi^* (M-M’) (n=42) and *PI3K^CAAX^*; *Dronc^RNAi^* (N-N’) (n=43). **(O-U)** *CHIZ>mGFP* (green) driven experimental sets [*Akt^RNAi^* (P) (n=52), *PI3K^DN^*(Q) (n=30), *InR^RNAi^* (R) (n=45)] show less Draper (red) compared to control, *CHIZ>mGFP/+* (O) (n=107); *CHIZ>mGFP* driven *PI3K^CAAX^* (S) (n=34) have significantly high Draper (red) and rescued in *PI3K^CAAX^; Drice^RNAi^* (T) (n=19) and *PI3K^CAAX^*; *Dronc^RNAi^* (U) (n=26). **(V)** Quantification of the number of Dcp-1 positive cells: control, *CHIZ>mGFP/+* (n=43) and *Akt^RNAi^* (n=50); control *CHIZ>mGFP/+* (n=32) and *PI3K^DN^* (n=37), *InR^RNAi^* (n=22) show significantly less Dcp-1 positive cells, control *CHIZ>mGFP/+* (n=30) with *PI3K^CAAX^* (n=29) show significantly high Dcp-1 positive cells and rescued in *PI3K^CAAX^; Drice^RNAi^*(n=42) and *PI3K^CAAX^*; *Dronc^RNAi^* (n=43). **(W)** Quantification of the number of γH2Av positive cells: control, *CHIZ>mGFP/+* (n=43) and *Akt^RNAi^* (n=47); control *CHIZ>mGFP/+* (n=40), *PI3K^DN^* (n=48), and *InR^RNAi^* (n=36) show significantly less γH2Av positive cells, control *CHIZ>mGFP/+* (n=30) with *PI3K^CAAX^* (n=29) show significantly high γH2Av positive cells and rescued in *PI3K^CAAX^; Drice^RNAi^* (n=42) and *PI3K^CAAX^*; *Dronc^RNAi^* (n=43). **(X)** Quantification of the Draper volume in control, *CHIZ>mGFP/+* (46) and *Akt^RNAi^* (n=52); control *CHIZ>mGFP/+* (n=35) and *PI3K^DN^* (n=30), *InR^RNAi^* (n=45) show significantly less Draper expression; control *CHIZ>mGFP/+* (n=26) with *PI3K^CAAX^* (n=34) show significantly high Draper expression and rescued in *PI3K^CAAX^; Drice^RNAi^* (n=19) and *PI3K^CAAX^*; *Dronc^RNAi^* (n=26). All images show a 25µm scale bar, with maximum intensity projections of the middle third optical section of wandering third instar lymph gland lobes. Nuclei stained with DAPI (blue). Control sets are different for groups of experimental sets in the graph (V, W, and X) because experimental sets are done on different days. Lymph glands boundary demarcated by white dotted line for clarity. ***P < 0.001, ****P < 0.0001 Error bars, mean ± SD. All images are representative of 3 or more independent biological experiments.

Next, we investigated if overactivation of PI3K/Akt signaling increases caspase activity and macrophage differentiation. A constitutively activated PI3K kinase (*UAS-PI3K^CAAX^*)^65^ expressed in the intermediate progenitors resulted in high caspase activity in all the CHIZ^+^ cells in the early third instar lymph gland (**Figures S4L-S4L’**). While the wandering third instar lymph glands showed a significant increase in Dcp-1-positive cells (**Figures 4A-4A’, 4E-4E’,** quantification showed in **4V**) and γH2Av-positive cell numbers (**Figures 4H-4H’, 4L-4L’,** quantification showed in **4W**). Also, the size of the lymph gland and CHIZ^+^ cell numbers increased significantly (quantification shown in **Figures S4C** and **S4D**). To test if the increase in caspase activity and DDR is a cell type-specific role of activated Akt or a direct response of a constitutively activated PI3K kinase expression in intermediate progenitors, we expressed *PI3K^CAAX^* and *Akt^RNAi^* in progenitor cells using the *dome^MESO^-Gal4* driver. Remarkably, in the *PI3K^CAAX^* background most lymph glands fell apart at the wandering third instar stage, the Dcp-1 positive and γH2Av positive cells were present in high numbers only in the differentiating zone instead in the core progenitor zone (**Figures S4N-S4O**), whereas *Akt^RNAi^* background showed smaller lymph glands with fewer γH2Av positive cells (**Figures S4N, S4P**). To determine if the increased caspase activity in the intermediate progenitors following *PI3K^CAAX^* overexpression affected macrophage differentiation, we performed Draper and P1 staining. We observed significantly high numbers of macrophages (**Figures 4O, 4S, S4H, S4I**, quantification in **4X, S4K**).

We found positive immunostaining for phosphorylated Akt (p-Akt) in the entire third instar lymph gland in control *CHIZ>mGFP* genotype, with less intense staining in the differentiated zone (**Figure S4Q-4Q’, S4T-S4T’**). Akt depleted intermediate progenitors (*CHIZ>mGFP>Akt^RNAi^*) resulted in decreased p-Akt (**Figures S4R-S4R’**, **S4U-S4U’**). However, p-Akt staining in *PI3K^CAAX^* overexpression using *CHIZ-GAL4*, which resulted in dramatically high p-Akt in CHIZ^+^ cells (**Figures S4S-4S’**, **S4V-S4V’**). These results confirm the presence of active Akt signaling in the intermediate progenitors and its role in macrophage differentiation.

Furthermore, depletion of caspases using *CHIZ-GAL4* driven *Drice^RNAi^* or *Dronc^RNAi^* in *PI3K^CAAX^*overexpression background lymph glands significantly reduced the number of Dcp-1 (**Figures 4A-4A’, 4E-4G’,** quantification in **4V**), γH2Av positive cells (**Figures 4H-4H’, 4L-4N’,** quantification in **4W**), Draper staining (**Figures 4O, 4S-4U,** quantification in **4X**) and lymph gland size (**Figures S4C, S4D**). These results thus clearly suggest that the InR-mediated PI3K/Akt-mediated signaling is upstream of apoptotic signaling involved in macrophage differentiation. However, the possibility of another receptor tyrosine kinase (e.g., Pvr, EGFR) playing a partially redundant role cannot be ruled out.

### Caspase-activated DNA breaks regulated by PI3K/Akt-mediated Ask1 attenuation

ROS-mediated Apoptosis signal-regulating kinase 1 (Ask1) activation promotes c-Jun N-terminal Kinase (JNK) signaling activation and apoptosis^66–68^. Surprisingly, the lymph gland differentiation is linked with ROS-mediated JNK activity^34^. Therefore, we hypothesized that Ask1 may activate caspase signaling in intermediate progenitors. Knockdown of Ask1 in intermediate progenitors (*CHIZ>mGFP>Ask1^RNAi^*) severely reduced immunostaining for extracellular protein matrix metalloprotease 1 (MMP1), a known JNK downstream reporter^69^ (**Figures 5A-5B’**) and Dcp-1 positive cell number significantly declined (**Figures 5G-5H’,** quantification in **5J**). In *Ask1^RNAi^*backgrounds DDR was significantly decreased (in two different *RNAi* lines, **Figures 5K-5L’, S5A-S5B’,** quantification in **5N**) along with severely affected macrophage differentiation (**Figures 5O-5P**, quantification in **5R**). This is consistent with previous reports that active JNK signaling contributes to progenitor differentiation^34, 70^. InR/PI3K/Akt signaling phosphorylate active p-Thr Ask1 on the N-terminal Ser83 residue attenuates Ask1 activity, resulting in a low level of MAP kinase activity^68^ that might activate a sublethal level of caspases. To test, we expressed a serine to alanine mutated Ask1 (*UAS-Ask1^S83A^*)^68^, which phenocopied the Ask1 knockdown phenotype of MMP1 staining (**Figures 5A-5A’, 5C-5C’**) and Dcp-1 positive cells reduced in number (**Figures 5G, 5I** quantification in **5J**). Expression of *Ask1^S83A^*also significantly reduced the number of γH2Av positive cells (**Figures 5K-K’**, **5M-M’**, quantification in **5N**) and Draper level (**Figures 5O, 5Q** quantification in **5R**). These findings indicate that *Ask1^S83A^*may act as a dominant negative allele^68^. Additionally, expression of *InR^RNAi^*, *Akt^RNAi^*, and *PI3K^DN^* in the intermediate progenitors (*CHIZ-Gal4*) also showed a dramatically reduced MMP1 staining as an indication of low JNK activity^69^ (**Figure 5A-5A’, 5D-5F’**). These data suggest that Akt signaling regulates Ask1-mediated JNK activation. Since ROS also regulates JNK activity in the lymph gland^34^, we used transgenic gstD-GFP as a ROS reporter^71^ and found γH2Av-positive cells have low gstD-GFP (**Figures S5C-S5C”**) compared to progenitors. Overall, our results indicate that lower ROS activity and Akt signaling promote an attenuated form of Ask1 (p-Ser83) rather than p-Thr Ask1 to control caspase activity at sublethal levels needed for macrophage differentiation.

**Figure 5.**
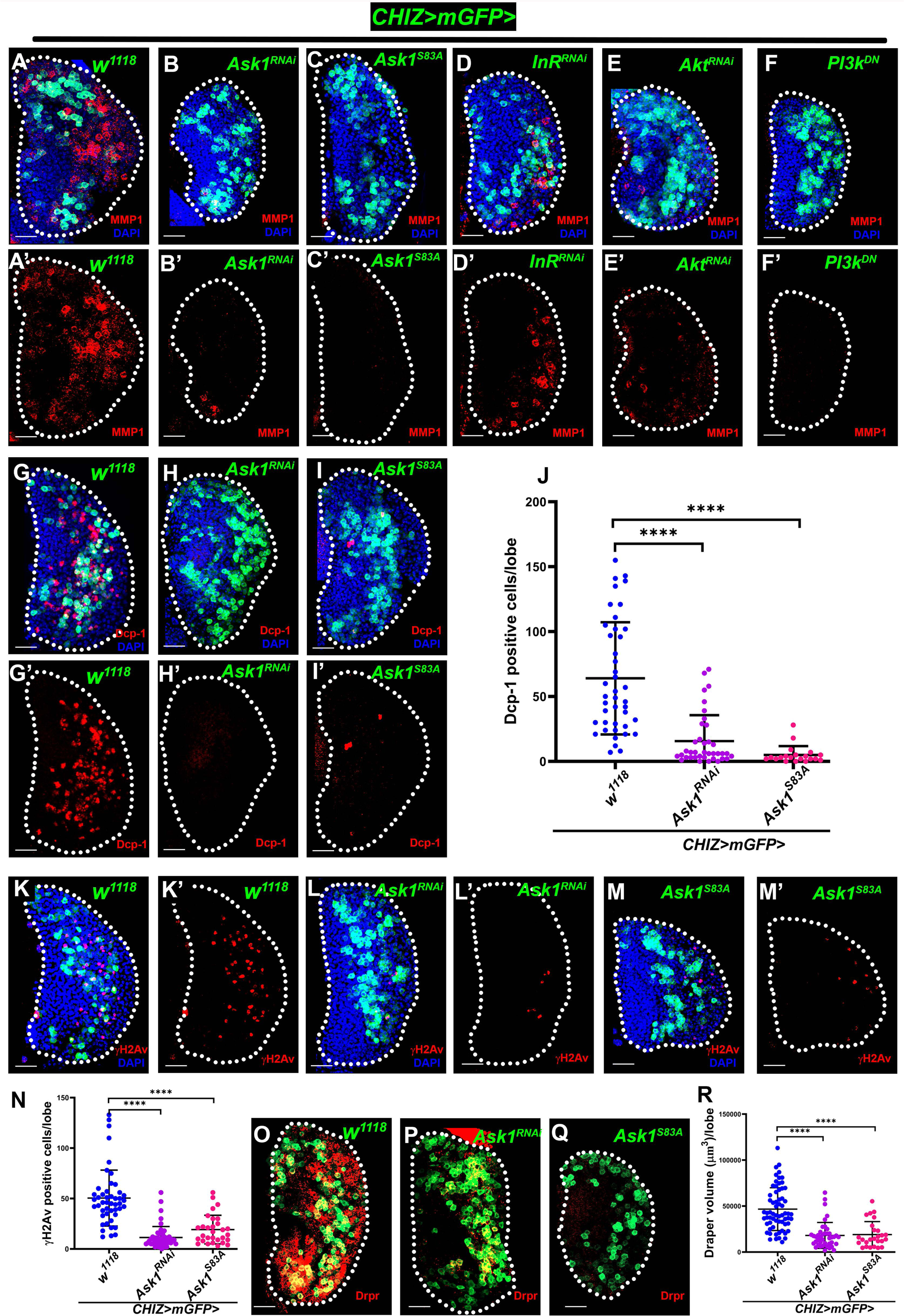
Caspase activity regulated by attenuated Ask1. **(A-F’)** JNK signaling activity in *CHIZ>mGFP* (green) driven *Ask1^RNAi^* (n=25) (B-B’), *Ask1^S83A^* (n=12) (C-C’), *InR^RNAi^* (n= 24) (D-D’), *Akt^RNAi^* (n=38) (E-E’) and *PI3K^DN^* (n=30) (F-F’) using MMP1 staining (red) severely reduces compared to control, *CHIZ>mGFP/+* (n=70) (A). **(G-I’)** *CHIZ>mGFP* driven *Ask1^RNAi^*(n=38) (H-H’) and *Ask1^S83A^* (n=20) (I-I’) show Dcp-1 positive cells (red) significantly decrease as compared to control, *CHIZ>mGFP/+* (n=42) (G-G’). **(J)** Quantification of the number of Dcp-1 positive lymph gland cells represented in (G-I’). **(K-M’)** *CHIZ>mGFP* driven *Ask1^RNAi^* (n=50) (L-L’) and *Ask1^S83A^*(n=30) (M-M’) show γH2Av positive cells (red) severely decrease as compared to control, *CHIZ>mGFP/+* (n=47) (K-K’). **(N)** Quantification of the number of γH2Av positive cells of lymph gland represented in (K-M’). **(O-Q’)** Draper staining (red) in control, *CHIZ>mGFP/+* (n=64) (O-O’), and in *CHIZ>mGFP* driven *Ask1^RNAi^*(n=41) (P-P’) and *Ask1^S83A^*(n=26) (Q-Q’) decrease significantly. **(R)** Quantification of the Draper volume of lymph gland represented in (O-Q’). All images are from wandering third instar lymph gland lobe with maximum-intensity projections of the middle third optical sections, scale bar is 25µm. Nuclei stained with DAPI (blue). Lymph glands boundary demarcated by white dotted line for clarity. ****P < 0.0001. Error bars, mean ± SD. All images are representative of 3 or more independent biological experiments. See also Figure S5.

### Caspase Activated DNase (CAD) induces DNA breaks required for macrophage differentiation

Among the diverse roles of caspases in myeloid-type progenitors, we explored the caspase-activated proteins that cause DNA breaks. Caspase-Activated DNase (CAD) is known to cause DNA breaks upon caspase activity, which cleaves its inhibitor ICAD (Inhibitor Caspase-Activated DNase). Free CAD dimerizes and functions as DNase, which causes DNA fragmentation during apoptosis^72, 73^. The CAD/ICAD proteins have not been extensively studied in *Drosophila*. The DNA fragmentation factor-related protein 1 (Drep1) is the ICAD homolog, and Drep4 is the CAD homolog in *Drosophila*^74–76^. We examined if *Drosophila* CAD/ICAD is responsible for DNA breaks in the lymph gland since we found caspase and DDR activity in the same cells (**Figure 2D**). We used lymph gland intermediate progenitor driver *CHIZ>mGFP* and progenitor driver *dome^MESO^-Gal4, UAS-2xEGFP* to knockdown *Drosophila* ICAD (*Drep1^RNAi^*) and CAD (*Drep4^RNAi^*) by multiple RNAi lines. The knockdown of ICAD and CAD in intermediate progenitors caused significantly fewer γH2Av-positive cells in the lymph gland (**Figures 6A-6C’**, **S6A-S6B’,** quantification showed in **6G**). Also, macrophage differentiation was significantly reduced, marked by Draper (**Figures 6D-6F**, quantitation in **6H**) and P1 (**Figures 6I-6K**, quantitation in **6L**). ICAD and CAD depletion in progenitors (*dome^MESO^>Drep1^RNAi^* or *Drep4^RNAi^*), also reduced γH2Av-positive cells (**Figures S6E-S6G**).

**Figure 6.**
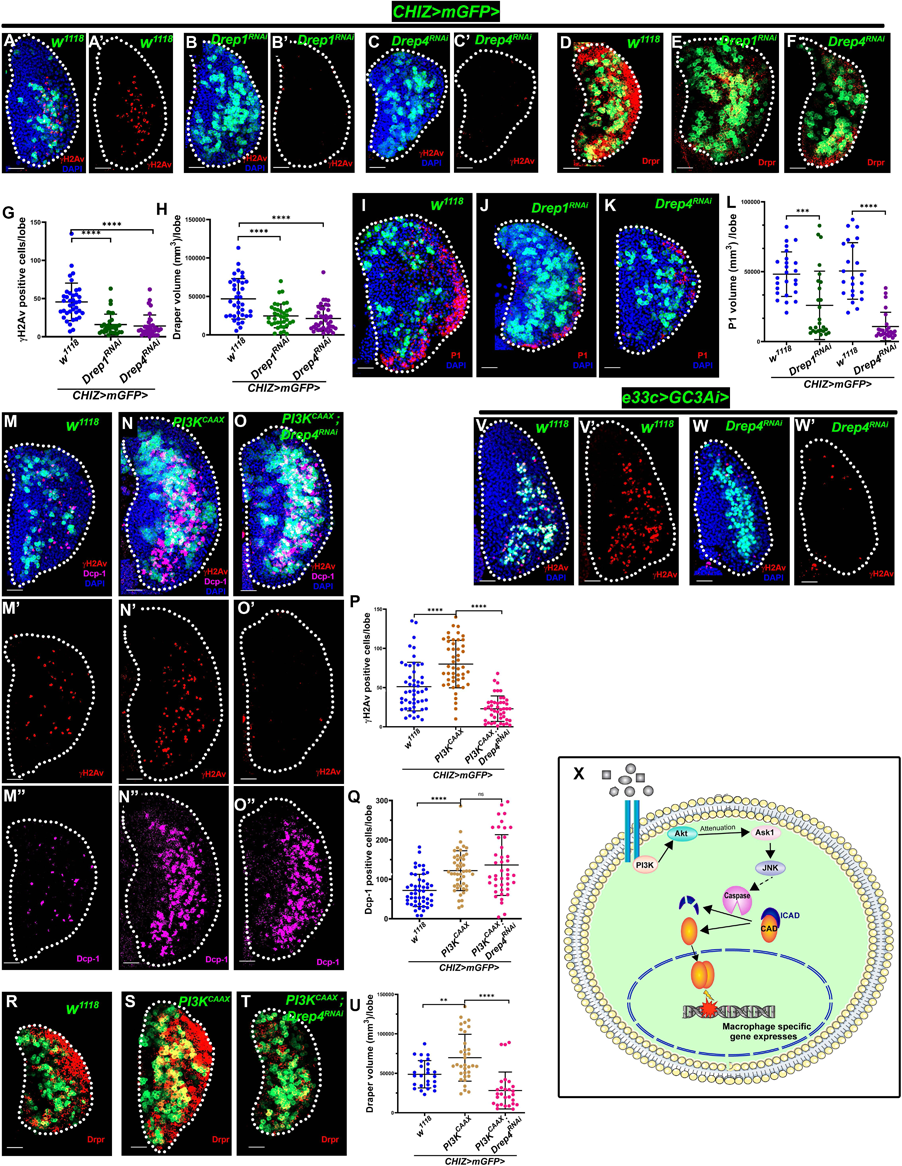
Caspase Activated DNase (CAD) induces DNA breaks required for macrophage differentiation. **(A-C’)** Depletion of *Drosophila* ICAD [(*CHIZ>mGFP; UAS*-*Drep1^RNAi^*) (n=34)] (B-B’), and CAD [*CHIZ>mGFP*; *UAS-Drep4^RNAi^*(n=35)] (C-C’) in the intermediate progenitors (green) lead to significantly less γH2Av positive cells (red) in the lymph glands compared to control, *CHIZ>mGFP/+* (n=38) (A-A’). **(D-F)** Phagocytic receptor marker Draper staining (red) in depletion of ICAD [*CHIZ>mGFP UAS*-*Drep1^RNAi^*(n=38)] (E), and CAD [*CHIZ>mGFP; UAS-Drep4^RNAi^*(n=40)] (F) cause significantly less Draper in the lymph gland compared to control, *CHIZ>mGFP/+* (n=37) (D). **(G)** Quantification of γH2Av positive cells of lymph gland represented in (A-C’). **(H)** Quantification of Draper volume of lymph gland represented in (D-F). **(I-K)** Macrophage marker P1 staining (red) with DAPI in depletion of ICAD [*CHIZ>mGFP UAS*-*Drep1^RNAi^*(n=26)] (J), and CAD [*CHIZ>mGFP; UAS-Drep4^RNAi^*(n=26)] (K) causes significantly less Draper in the lymph gland compared to control, *CHIZ>mGFP/+* (n=23) (I) GFP cell marks intermediate progenitors (green). **(L)** Quantification of P1 volume of lymph gland represented in (I-K). **(M-O’’)** Overexpression of PI3K active form in intermediate progenitors [*CHIZ>mGFP*; *UAS*-*PI3K^CAAX^* (n=48)] (N-N’’) causes significantly high γH2Av and Dcp-1 positive cells (red) compared to control, *CHIZ>mGFP/+* (n=50) (M-M’’), and depletion of CAD [CHIZ>mGFP*; UAS*-*PI3K^CAAX^; UAS-Drep4^RNAi^* (n=45 for γH2Av and n=42 for Dcp-1 staining)] rescue high number of γH2Av positive cells (N’ and O’), however number of Dcp-1 positive cells (N” and O”) remain same as *PI3K^CAAX^* expression background. **(P)** Quantification of γH2Av positive cells of lymph gland represented in (M’-O’). **(Q)** Quantification of Dcp-1 positive cells of lymph gland represented in (M”-O’’). **(R-T)** Overexpression of PI3K active form in intermediate progenitors [*CHIZ>mGFP*; *UAS*-*PI3K^CAAX^* (n=34)] (S) causes high macrophage marker Draper (red) and large lymph gland compared to control, *CHIZ>mGFP/+* (n=26) (R), and depletion of CAD [*CHIZ>mGFP; UAS PI3K^CAAX^; UAS-Drep4^RNAi^* (n=28)] (T) rescue high Draper in *PI3K^CAAX^* expression background (S). **(U)** Quantification of Draper staining volume of lymph gland represented in (R-T). **(V-W’)** Loss of CAD in the lymph gland (*UAS*-*GC3Ai/UAS-Drep4^RNAi^; e33c-Gal4/+*) (n=24) (W-W’) show less γH2Av positive cells (red) but caspase active (*GC3Ai*) cells remain unchanged compared to control, (*UAS*-*GC3Ai/+; e33c-Gal4/+*) (n=25) (V-V’). **(X)** For macrophage differentiation, a model showing InR/PI3K/Akt/Ask1 mediated caspase activity and CAD-mediated DNA breaks. All images are from wandering third instar larval lymph gland lobe. Scale bar is 25µm, and images are maximum intensity projections of the middle third optical section. Nuclei stained with DAPI (blue). Lymph glands boundary demarcated by white dotted line for clarity. **P < 0.001, ****P < 0.0001, ns-not significant Error bars, mean ± SD. All images are representative of 3 or more independent biological experiments.

Since antibodies against *Drosophila* CAD/ICAD are not available, we used *Drep4 T2A-Gal4* line^77^, a CRIMIC line that likely recapitulates *Drep4* (*CAD*) gene expression patterns to drive *UAS-mRFP*. Most lymph gland cells are *Drep4>mRFP*+ (**Figure S6H**). We used quantitative RT-PCR to assess the transcript level of *Drep1/Drep4* (*ICAD/CAD*) in larval tissues and the efficiency of RNAi lines. Both CAD and ICAD genes expressed in larval tissue, and RNAi lines effectively reduced their transcript levels (**Figures S6I-S6J**).

DNaseII and Endonuclease G (Endo G) also function with alternative apoptotic signaling and contribute to DNA breaks similar to CAD^78, 79^. To assess the role of DNaseII and Endo G in blood progenitor differentiation, we used γH2Av immunostaining in the homozygous *DNaseII^lo^* hypomorphic allele and Endo G mutant (*EndoG^MB071^*^50^)^79^. The γH2Av positive cell numbers were unaffected in both homozygous mutants (**Figure S6K-S6M**). Thus, the DNaseII and Endo G were not involved in the DNA damage-mediated macrophage differentiation.

To determine if knockdown of the *Drosophila* CAD/ICAD complex in *PI3K^CAAX^* overexpression background alleviates the caspase activity and DNA damage, we used the *CHIZ-GAL4* driver to knock down CAD (*Drep4)* in *PI3K^CAAX^* overexpression background. This intervention severely reduced the number of γH2Av-positive cells than in the control and overexpression backgrounds (**Figures 6M-6O’**, quantification showed in **6P**). However, it is noteworthy that the Dcp-1 positive cells remained high similar to overexpression of *PI3K^CAAX^*(**Figures 6M”-6O”**, quantification showed in **6P**), and lymph gland size (by measuring DAPI stained cells volume) and CHIZ^+^ positive cells unaffected (**Figures S6C-S6D**) when compared with the PI3K^CAAX^ overexpression background. Also, macrophage differentiation was noticeably decreased following the knockdown of CAD with *PI3K^CAAX^*overexpression background (**Figures 6R-6T**, quantification showed in **6U**). Further, depleting CAD (Drep4) in the whole lymph glands (*e33c>GC3Ai*>*Drep4^RNAi^*) did not affect the caspase activity but mostly reduced γH2Av positive cells (**Figure 6V-6W’**), which indicates that CAD causes DNA breaks in the lymph gland during macrophage development. Thus, these findings show that the differentiation of macrophages relies on CAD/ICAD-mediated DNA damage induced by InR/PI3K/Akt signaling in the lymph gland (**Figure 6X**).

### Autophagy-mediated survival of the phagocytic macrophages

The *Drosophila* Autophagy-related 6 (Atg6) mutant has dysplastic lymph glands with no macrophage differentiation, while the Atg1 mutant has circulating macrophage spreading defects^80, 81^. Autophagy induction is also critical for monocytes to macrophage differentiation and survival^12, 82, 83^. As autophagy is already known to be involved in macrophage differentiation, we emphasize checking whether autophagy occurs in the γH2Av-positive cells in the lymph gland. We first expressed the autophagy reporter *UAS-GFP-mCherry-Atg8a*^84^ with the *e33c-Gal4* driver to detect autophagy status in lymph gland cells. We observed that mCherry-Atg8a autophagic puncta were abundant in intermediate progenitors and differentiated zone (**Figure 7A-7A’, S7A-A’**). γH2Av-positive lymph gland cells had large mCherry-Atg8a autophagosomes puncta (**Figures 7B-7B’, S7B**). We used the Anti-GABARAP (anti-Atg8a) antibody^85, 86^ (Abcam, ab109364) to detect endogenous *Drosophila* Atg8a and autophagosomes. Remarkably, lymph gland γH2Av-positive cells showed a high number of autophagic puncta (**Figures 7C-7D’**). We also marked acidic compartments of live cells that grow during autophagy with LysoTracker Red^85^. LysoTracker-positive cells are visible in the intermediate zone of lymph glands (**Figures 7E-7F’, S7C-S7C’**) and control fat body tissue (**Figure S7D-S7D’**). Besides, ROS reporter gstD-GFP-positive cells with low GFP showed mCherry-Atg8a autophagy puncta (**Figures S7E-S7E”’**), this indicates autophagy requires in the survival of γH2Av positive cells. Based on published studies^80–82, 87^ and our findings, autophagy appears to have a role in the survival of caspase-activated DNA breaks differentiating macrophages. We proposed a model for *Drosophila* myeloid-type progenitors to macrophage differentiation via developmental signaling-induced caspase-activated DNA breaks in **Figure 7G**.

**Figure 7.**
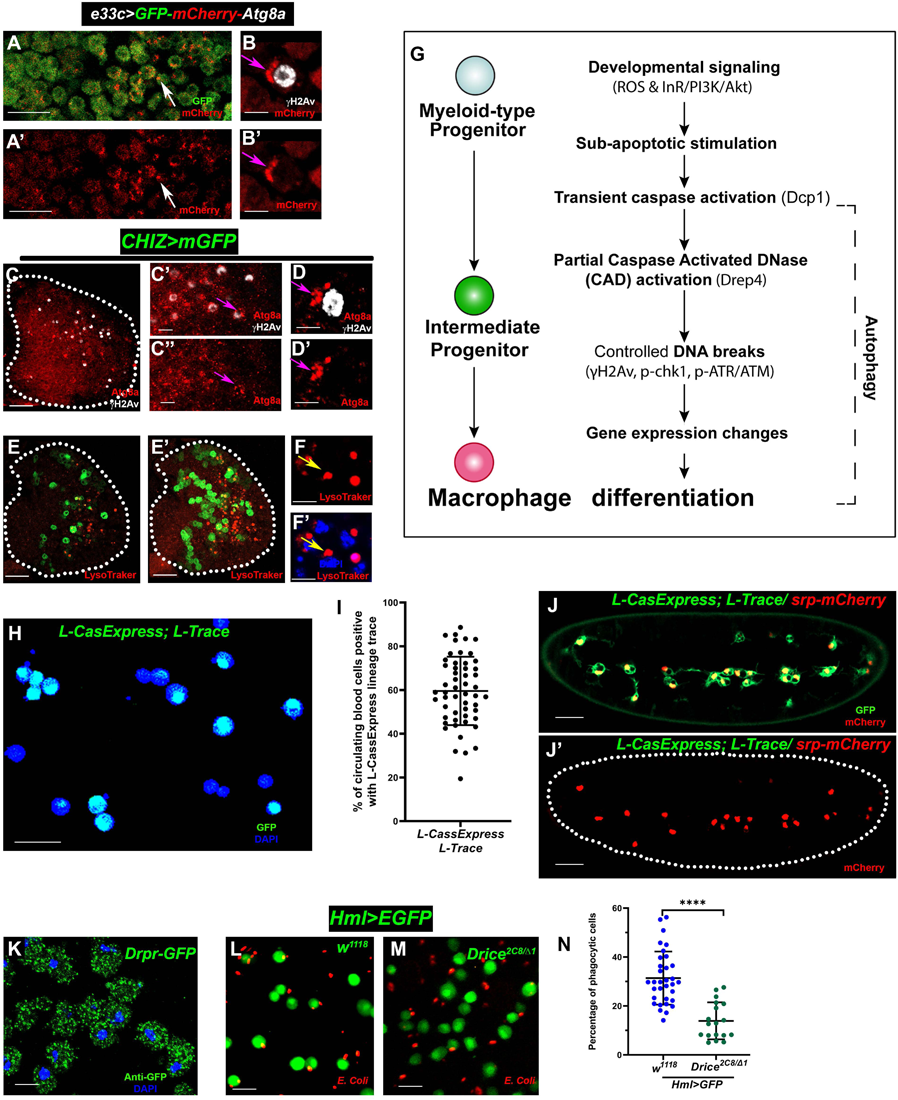
Developmental caspase-activated DNA breakage and autophagy regulate macrophage differentiation. **(A-A’)** *e33c-Gal4, UAS-GFP-mCherry-Atg8a* expression in lymph gland shows *Atg8a-mCherry* puncta marked by arrow(A-A’) in the differentiated zone, also see lymph gland in Figure S7. (B-B’) High magnification of *e33c-Gal4, UAS-GFP-mCherry-Atg8a* expression of lymph gland shows *Atg8a-mCherry* puncta in γH2Av (grey) positive cell marked by arrow. **(C-D’)** Antibody against Atg8a (red) co-staining with γH2Av (grey) in control *CHIZ>mGFP/+* (without GFP) (C) and inset images show colocalization of Atg8a and γH2Av in the same cell (C’-C’’), also show in high magnification (D-D’) marked by an arrow. **(E-F’)** LysoTracker-positive cells are visible in the intermediate zone of live lymph glands by LysoTracker staining (red) in control CHIZ>mGFP/+ (green), showing a single optical section (E) and middle third projection (E’). It is also shown in high magnification (F-F’) marked by the arrow. **(G)** Schematic shows that the mechanism of myeloid-type progenitor to macrophage differentiation through intermediate progenitor requires transient caspase activation and CAD-mediated DNA breaks. **(H)** Circulating blood cells of third instar larvae *L-CasExpress L-Trace* (n=53) showing caspase lineage activity (GFP) (green). **(I)** Quantification of the represented image in (H) shows 60% caspase lineage-positive circulating cells. **(J-J’)** Embryonic macrophages (stage 13) marked by *srp-mCherry* (red) are caspase lineage (*L-CasExpress L-Trace*) GFP (green) (J-J’) positive. **(K)** Circulating blood cells of third instar *Draper-GFP* larvae show Draper (green) punctate staining. **(L-M)** Phagocytosis of RFP tagged *E. coli* by 3^rd^ instar circulating macrophages in control *Hml^Δ^-Gal4, UAS-2xEGFP*/+ (green) (L) and mutant *Hml^Δ^-Gal4, UAS-2xEGFP /+; Drice^2c8/Δ1^*(n=18) (M) show less phagocytic efficiency. **(O)** Quantification of the percentage of phagocytic circulating macrophages represented in (L-M). All the lymph gland images shown from the wandering third instar lymph gland, except J-J’ are stage 13 embryo. Scale bars: 25µm in all images except 10µm for images (C’-C’’, and K-M), 5µm for images (B-B’, D-D’, and F-F’). All images are single optical sections except image E’, which is the maximum intensity projection of the middle third optical section. Nuclei stained with DAPI (blue). Lymph glands boundary demarcated by white dotted line for clarity. ****P < 0.0001, ns-not significant Error bars, mean ± SD. All images are representative of 3 or more independent biological experiments.

### Embryonic-origin macrophages require caspase activation for efficient phagocytoses

Like vertebrates, *Drosophila* hematopoiesis occurs during early embryogenesis. Embryonic macrophages disperse throughout the embryo, later populating the larval sessile and circulating blood cells^88^. Therefore, we first examined embryo-origin circulating blood cells from third-instar larvae to see whether the caspase lineage reporter (*L-CasExpress L-Trace*) can mark caspase-activated lineage cells with GFP^53^. Remarkably, 60% of circulating cells are caspase lineage-positive (**Figure 7H**, quantification showed in **7I**). Embryonic macrophages express homolog of GATA transcription factor serpent (srp)^89^. To find if macrophages in embryos also experienced executioner caspase activation marked with the caspase lineage reporter, we live-imaged embryos using *L-CasExpress L-Trace* and *CasExpress-Gal4; G-Trace^LTO^* in the *srp-mCherry*^89^ background, where *srp-mCherry* marked embryonic macrophages. Remarkably, macrophages were co-localized with caspase reporter L-*CasExpress*-GFP positive cells (**Figure 7J-7J’**), and also co-localized with CasExpress>GFP positive cells (**Figure S7F-S7G’** and **Movies S1-S2**). But a vast number of *L-caspase L-Trace* lineage positive cells are found in the dorsal closure region (**Figure S7H-S7H’** and **Movie S3**), at developmental stage 13, when many cells are known die and served as a control tissue for our experiments. Draper is a single-pass transmembrane receptor involved in phagocytosis and also for autophagy activation^90^. Thus, we monitored Draper expression using *Draper-GFP* line^91^, which highly expressed in embryonic-origin larval circulating blood cells (**Figure 7K**) and Draper antibody staining co-localizes with Draper-GFP in the lymph gland differentiated zone (**Figure S7I-S7I”**). Finally, we performed a phagocytic assay^92^ using fluorescently labeled *E. coli* in wandering third-instar circulating macrophages marked by *Hml^Δ^-Gal4, UAS-2xEGFP* and found a significant decrease in the phagocytic efficiency of bacteria in caspase mutant (*Drice*^*2c8*/Δ1^) macrophages (**Figure 7L-7M**, quantification showed in **7N**). Collectively, our data suggest that caspase-activated DNA breaks are essential for *Drosophila* phagocytic macrophage differentiation.

## Discussion

Macrophages are multifunctional phagocytic cells in the innate immune system that populate most tissues during early fetal development and maintain themselves independently of adult hematopoiesis through longevity and limited self-renewal^1,2,4^. However, the macrophage differentiation mechanisms remain unknown.

Here, we show that during the normal development of *Drosophila* macrophages in the larval lymph gland, apoptotic caspases are activated in the differentiating cells. A sublethal level of executioner caspase activation induces CAD, triggering DNA strand breaks in the differentiating macrophage. High ROS signals from the progenitors induce the apoptotic signal through Ask1/JNK signaling activation. We also find that InR/PI3K/Akt-mediated signaling activates apoptotic signaling in the intermediate progenitors. The Akt signaling contributes to the attenuation of Ask1 by phosphorylating it at Ser83 residue in differentiating macrophages. DNA damaged differentiating cells display autophagy activity that may use as a macrophage differentiation cum survival strategy. Furthermore, caspase activation is required in embryonic-origin macrophage development for efficient phagocytic activity. Therefore, our research using in vivo genetic analysis revealed a previously undiscovered and unique role that growth factors signaling, caspases, and DDR signals play in determining macrophage differentiation during normal development.

Many *Drosophila* cells show caspase activation to have non-lethal roles in development and differentiation, as shown by several labs^16, 47, 51, 56, 93–95^. Also, monocyte-to-macrophage differentiation requires CSF1-Akt-mediated caspase activation^96^. ROS has been shown to modulate caspase activation during *Drosophila* thorax fusion development^94^. The lymph gland progenitors have high ROS-mediated JNK signaling, which is critical for progenitor maintenance and differentiation^34^. Thus, we hypothesize that ROS plays a role in activating caspases in this system. Here, DDR-positive cells have low ROS, which explains that ROS might cause caspase activation, and by the time DDR is active, ROS becomes low as the intracellular environment also changes. We find that InR-mediated PI3K-Akt signaling has a dual role for autonomous apoptotic activation and control caspase activity, as reported^53^. However, the possibility of other signaling (e.g., Pvr^28^, EGFR, GABA-Calcium^30, 32^) playing a partially redundant role cannot be ruled out. We find that both initiator and effector caspase are required for Draper expression in the differentiated macrophages. Consistent with our data, embryonic hemocytes have low levels of Draper in the lack of *RHG* genes^97^. Our lineage-trace experiment for caspase-positive cells confirms that differentiated macrophages come through caspase-activation. Moreover, we demonstrate a substantial reduction in DDR-positive cells and phagocytic cells through various manipulations of the apoptotic signal in intermediate progenitors. Here, we reveal an unknown mechanism: developmental signal-mediated caspase activation during embryonic and larval macrophage differentiation.

Programmed DNA breaks coordinate gene expression changes without cell death in several types of cell differentiation^98^, but the signals that cause DNA damage were not addressed. Interestingly, single-cell transcriptomics on lymph glands revealed a group of cells (1.2%) called cluster X or GST-rich with unique genetics and enrichment of DDR, Myb, and cell cycle genes^70, 99^. These cells are most likely the CAD-mediated DNA-damaged cells here we report as the location and cell numbers in the lymph gland are comparable. Overall, this study revealed for the first time that caspase-mediated *Drosophila* CAD causes DNA breaks, which is essential for macrophage differentiation as depletion of CAD or ICAD in the lymph gland causes loss of phagocytic markers and DNA damage, but caspase activity is still seen. Also, CAD depletion rescues the PI3K active phenotypes except for caspase activity, which suggest InR/PI3K-mediated CAD activity is required for macrophage differentiation (**Figure 7G**).

How well levels and dynamics of executioner caspase predict cell survival vs. cell death remains unclear. However, using a cancer cell line model, it was showen that high caspase activity kill all cells, but low levels allow cells to survive^100^. Here, we show that Akt signaling attenuates Ask1 activity that controls caspase and CAD activity at sublethal level in the differentiating cells. The autophagy mechanism is used in various cells for survival^12^, including monocyte-to-macrophage differentiation^82, 83^ along with PI3K/Akt/Tor signaling involvement^101^. Here, γH2Av positive cells with high autophagy puncta explain that autophagy may operate as a survival and differentiation mechanism. Other mechanisms might also help survival after caspase activation^102, 103^. For example, in vitro studies of caspase-mediated skeletal muscle cell differentiation reported that nuclear pore complex trimming alters the intracellular environment^104^, and CAD-mediated DNA damage is repaired by base excision repair protein XRCC1, resulting in gene expression changes^41, 105^. Differential accessibility of transient CAD for DNA fragmentations helps cells survive due to their chromatin architecture^106, 107^.

Caspase/CAD causes DNA breaks around chromatin-modifying CCCTC-binding factor (CTCF) sites (chromatin insulators) to control gene expression by directly acting on a promoter or influencing promoter-enhancer interactions^108–110^. A study showed that the enrichment of insulator proteins at insulator sites rises after DNA damage by regulating γH2Av in *Drosophila*^111^. Moreover, mammalian macrophage functions require a set of transcriptional regulators accomplished by the tissue microenvironment in the tissue-specific macrophage chromatin landscape^5,6^. Our findings suggest that the survival of DNA-damaged progenitor cells may influence the specification of macrophage fate by modulating chromatin landscape and gene expression. Caspase/CAD-mediated DNA breaks in differentiating macrophages prepare them for various types of infection and injury in the future, activating macrophages as predicted in trained immunity^20, 97, 112^ and efficient tissue-specific functions^1,2,4^. Further research will determine whether this is also relevant to macrophages in higher organisms.

## Acknowledgments

We thank U. Banerjee, A. Bergmann, M. Miura, B. Hay, M. Suzanne, E. Wieschaus, M. Inamdar, D. Bohmann, I.K. Hariharan, F. Serras, I. Ando, T. Mukherjee for reagents; J. Barman, J. Biswas for preliminary data; S.C. Lakhotia and Cytogenetics laboratory members for valuable inputs and support. We acknowledge BDSC, VDRC for fly stocks, and DSHB for antibodies. This study was funded by the DBT/Wellcome Trust India Alliance Intermediate Fellowship (IA/I/20/1/504931), DBT-Ramalingaswami Fellowship (BT/RLF/Re-entry/08/2016), and Institute of Eminence Scheme, BHU to BCM, and CSIR fellowship to DM and DM.

## Author contributions

Conceptualization, B.C.M. and D.M.; Methodology, B.C.M. and D.M.; Investigation and analysis, D.M.; Visualization, D.M, G.R., and D. Mandal; Model preparation, G.R., B.C.M. and D.M; Writing, B.C.M. and D.M.; Funding acquisition and supervision, B.C.M.

## MATERIALS AND METHODS

### KEY RESOURCE TABLE

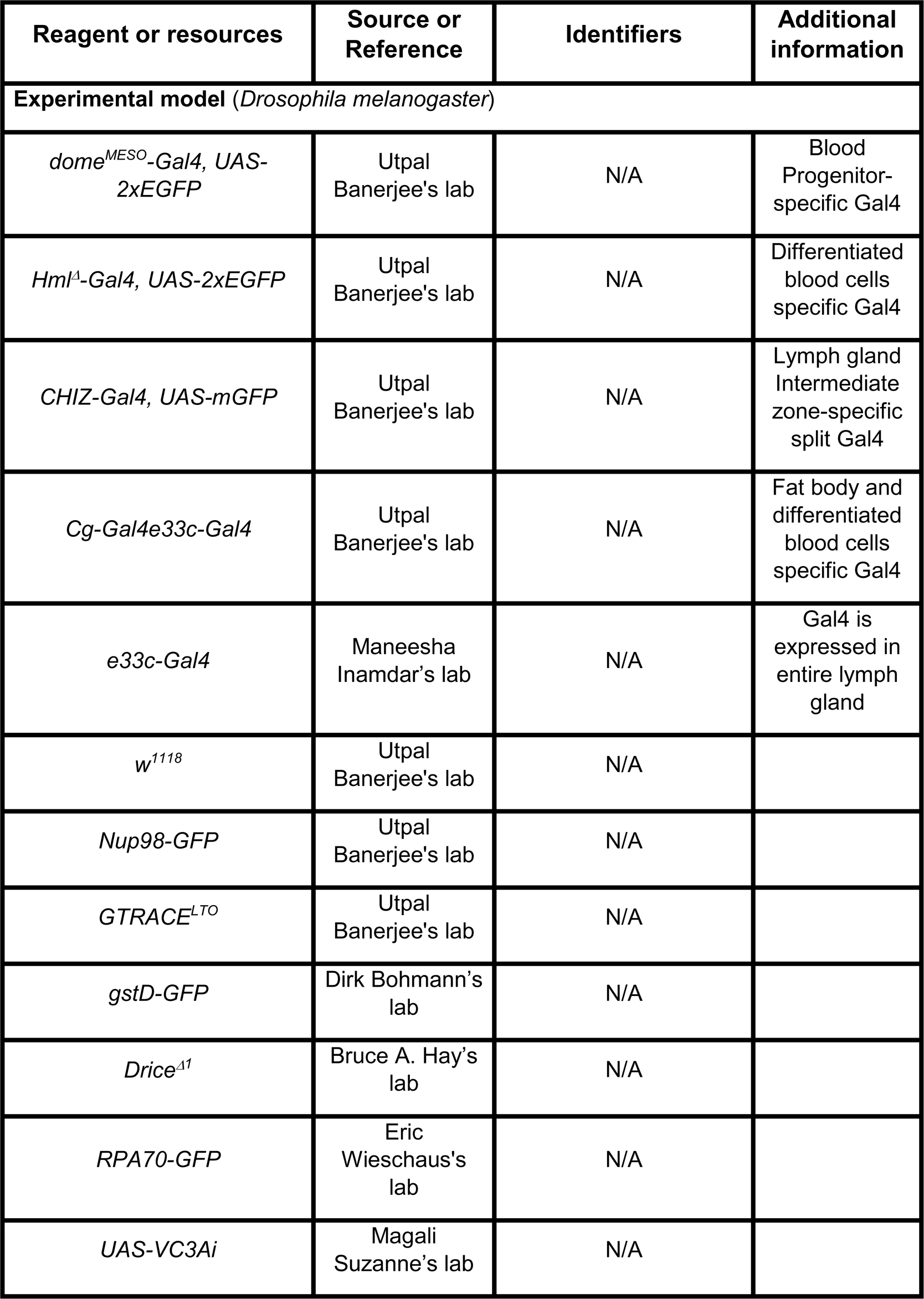
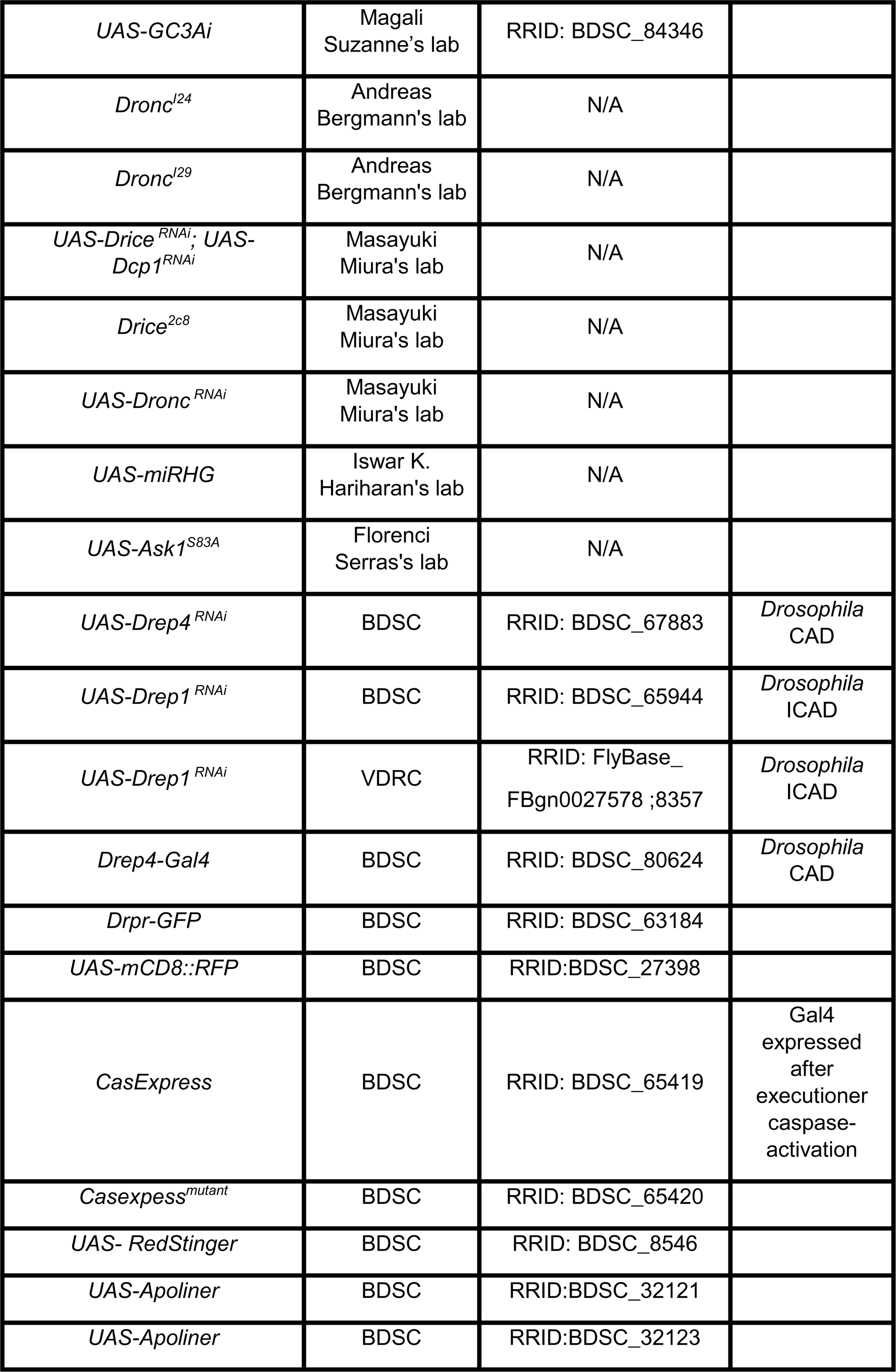
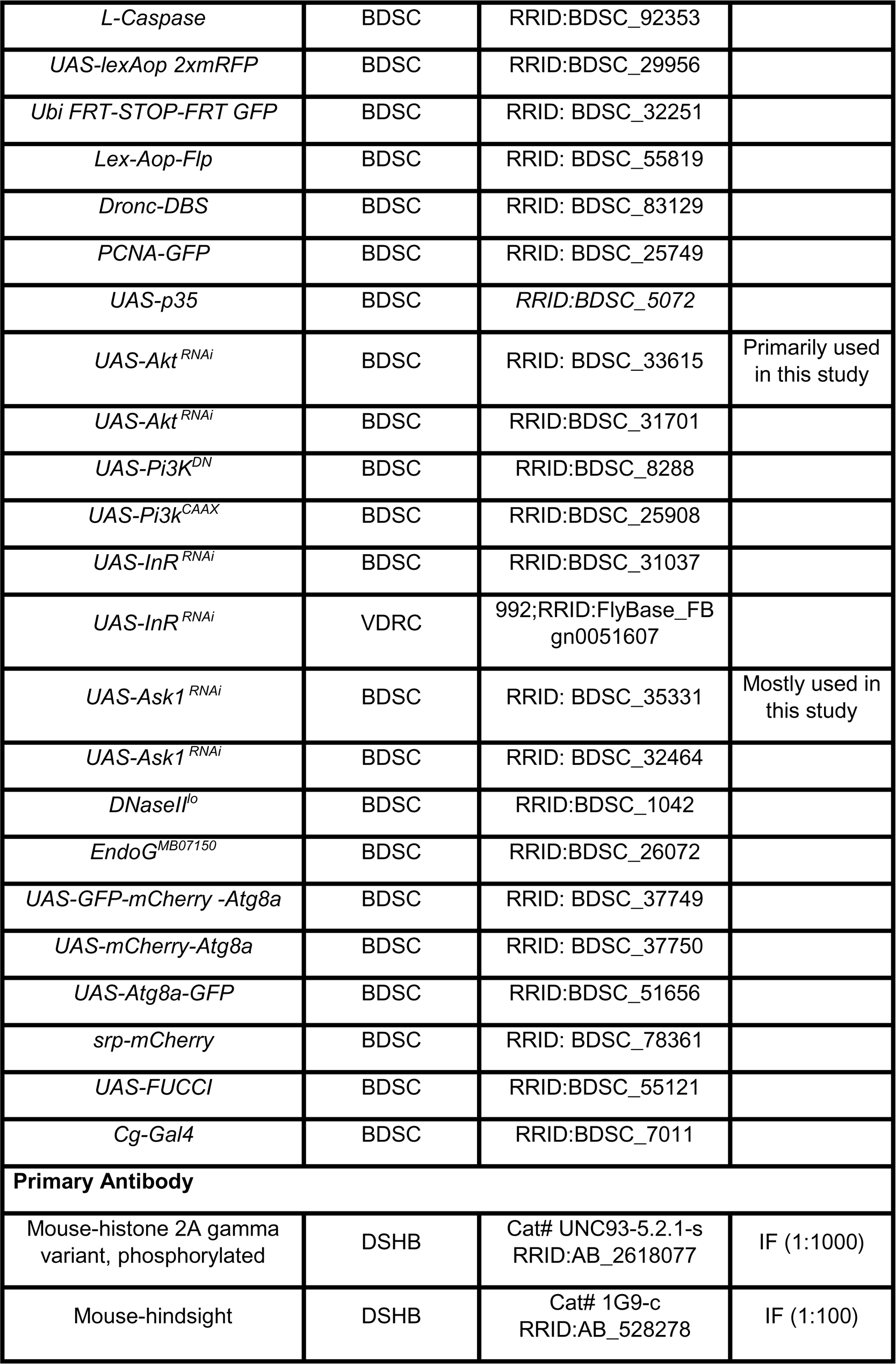
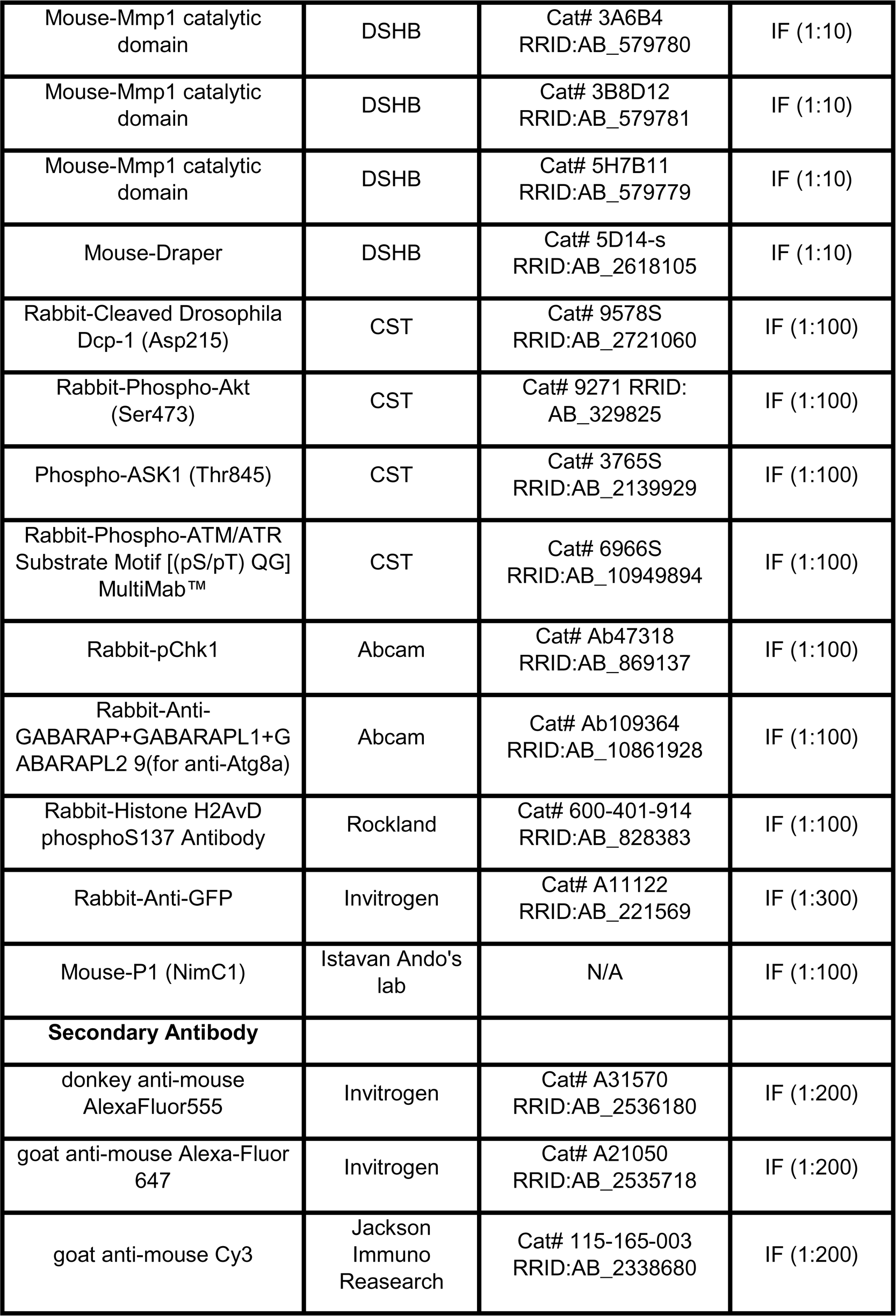
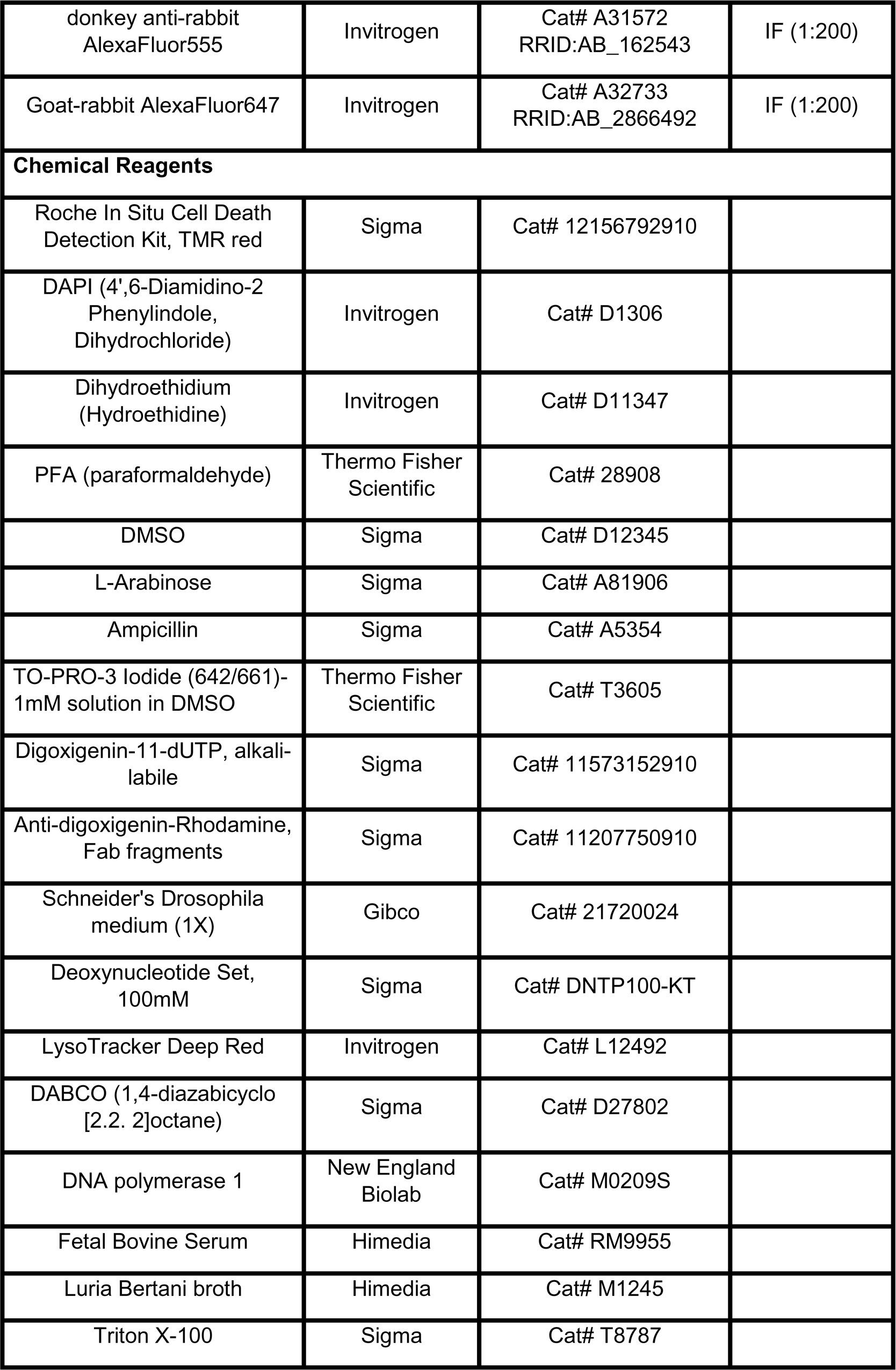
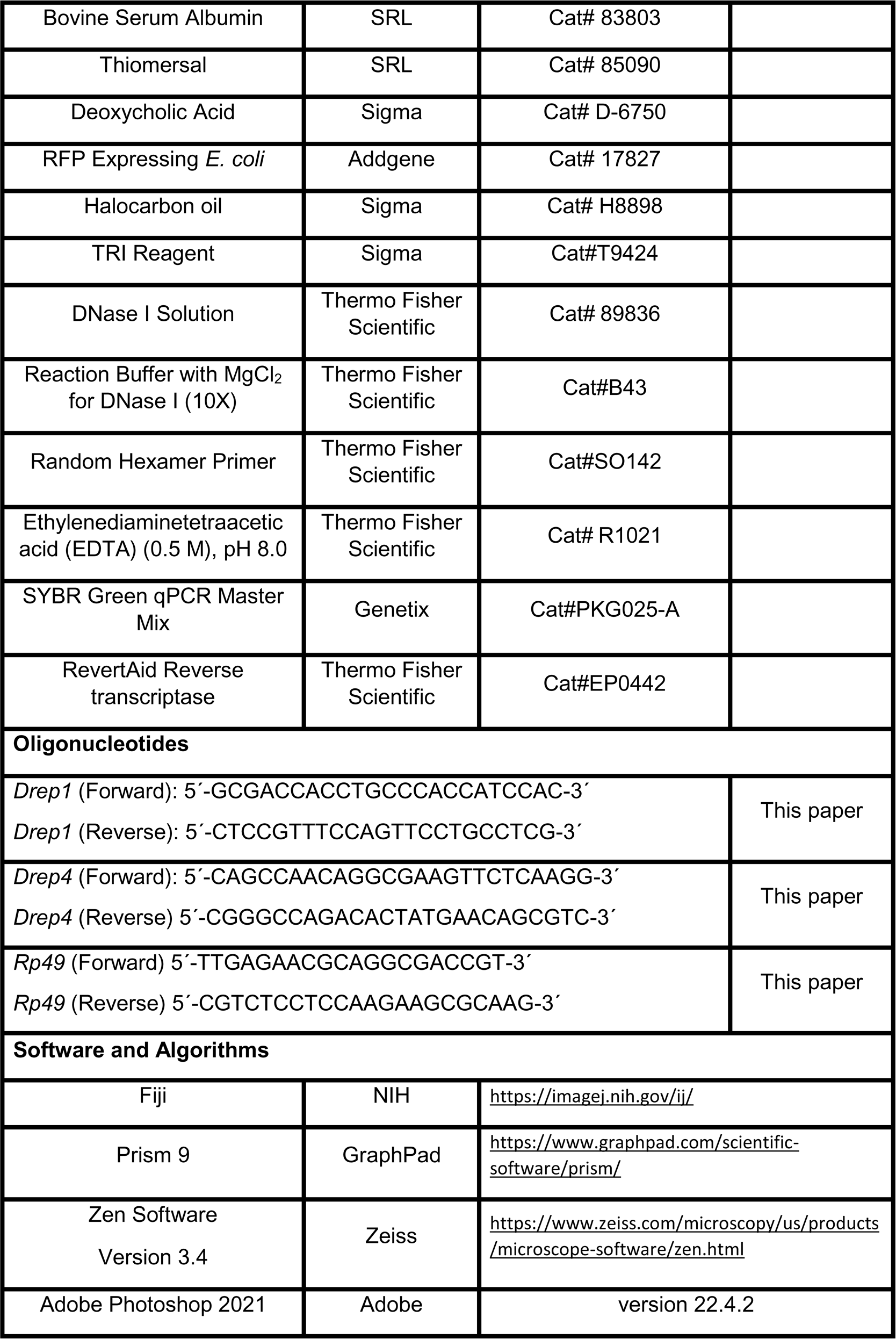
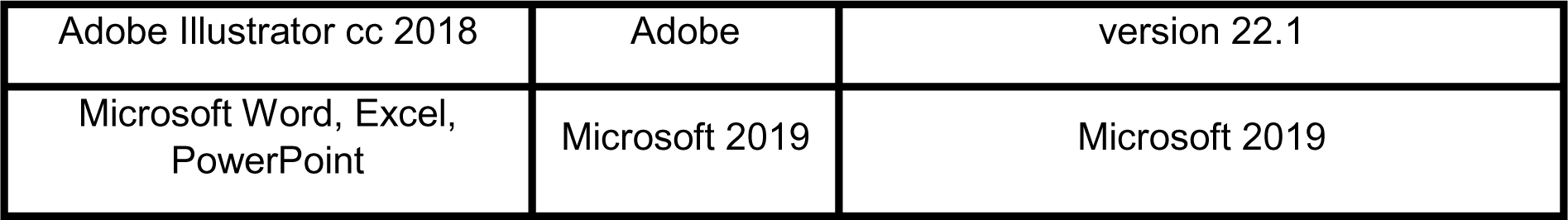

#### Lead contact

Further information and requests for resources and reagents should be directed to and will be fulfilled by the lead contact, Bama Charan Mondal.

#### Materials Availability

This study did not generate new reagents.

#### Data and Code Availability

The lead contact will share all data reported in this paper upon request. This study did not generate any unique datasets or code.

### EXPERIMENTAL MODEL AND SUBJECT DETAILS

#### *Drosophila* stocks and husbandry

*Drosophila* stocks were cultured using standard fly medium comprising 46g/L cornmeal, 45g/l sucrose, 18g/l yeast extract, 7g/l agar, supplemented with 3ml/L propionic acid, and 3g/l p-hydroxybenzoic acid methyl ester. All stocks were maintained at room temperature or 18°C, and genetic crosses using the GAL4/UAS system were maintained at 29°C on a 12 h light/ 12 h dark cycle. The following *Drosophila* stocks were used for this study: *CHIZ-GAL4 UAS-mCD8::GFP^39^, Hml^Δ^-Gal4 UAS-2xEGFP, dome^MESO^-Gal4 UAS-2xEGFP, Nup98-GFP, w*^111^*^8^, UAS-RedStinger* (BL8546), and *UAS-GTRACE^LTO^*(BL28282) were from Utpal Banerjee’s lab. The following fly lines were obtained from Bloomington *Drosophila* Stock Center (BDSC): *UAS-Akt^RNAi^* (BL33615 and BL31701), *UAS-Ask1^RNAi^* (BL35331 and BL32464), *UAS-Drep1^RNAi^* (BL65944), *UAS-Drep4 ^RNAi^*(BL67883), *UAS-InR^RNAi^* (BL31037), *UAS-GC3Ai* (BL84346), *srp-mCherry* (BL78361)^89^, *DnaseII^lo^* (BL1042), *Dronc-DBS* (BL83129), *Ubi-p63-(FRT-STOP-FRT Stinger*) (BL32250), *L-Caspase* (BL92353)^53^, *LexAop-Flp* (BL55820), *PCNA-GFP* (BL25749), *UAS-Apoliner* (BL32121 and BL32123), *EndoG*^*MB07150*^ (BL26072), *Drpr-GFP* (BL63184), *UAS-PI3K^DN^* (BL8288), *UAS-PI3K^CAAX^* (BL25908), *UAS-GFP-mCherry-Atg8a* (BL37749)*, UAS-Atg8a-GFP* (BL51656), *UAS-mCherry-Atg8a* (BL37750), *UAS-lexAop-2xmRFP* (BL29956), *UAS-p35* (BL5072), *CasExpress^mutant^* (BL65419), *CasExpress* (BL65420)^47^, *UAS-mCD8::RFP* (BL27398), *Drep4-Gal4* (BL80624)^77^, *UAS-FUCCI* (BL55121), *Cg-Gal4* (BL7011). Flies from Vienna Drosophila Stock Center: *UAS-Drep1^RNAi^*(v8357) and *UAS-InR ^RNAi^* (v992). The following stocks were kind gifts from different labs: *Dronc^I29^*, *Dronc^I24^* (Andreas Bergmann)^57^, *Drice^Δ1^* (Bruce A Hay)^56^*, Drice*^*2c8*^, UAS-Drice ^RNAi^; UAS-Dcp1 ^*RNAi*^ and *UAS Dronc ^RNAi^*(Masayuki Miura)^55,61^; *UAS-miRHG* (Iswar K. Hariharan)^60^; *UAS-Ask1^S83A^* (Florenci Serras)^68^; *RPA70-GFP* (Eric Wieschaus)^44^; *e33c-Gal4* (Maneesha Inamdar)^26^, *gstD-GFP* (Dirk Bohmann)^71^, *UAS-GC3Ai, UAS-VC3Ai* (Magali Suzanne)^48^.

#### *Drosophila* lymph gland dissection and immunostaining

Lymph glands (LGs) were dissected from wandering third instar larvae on a silicon dissecting plate. The head complex was isolated, comprising of lymph gland, brain, eye-antennal disc, and mouth hook, in chilled 1X PBS (phosphate buffer saline). The tissues were then immersed in a fixative solution, 4% paraformaldehyde (PFA, Thermo Fisher Scientific, Cat# 28908) in 1X PBS for 30 minutes and washed 3 times for 10 minutes each with wash buffer (0.3% Triton X-100 in 1X PBS). Samples were incubated with blocking solution (0.1% Triton X-100, 0.1% BSA, 10% FBS, 0.1% deoxycholate, 0.02% thiomersal) for 2 hours at room temperature (or in the case of Draper staining 24 hours at 4°C) and then incubated with primary antibody overnight at 4°C. Samples were washed with a wash buffer thrice, then incubated with a blocking solution for 2 hours at RT and incubated with a secondary antibody for 2 hours at RT. Following the incubation with the secondary antibody, the tissues were subjected to three washes in 0.3% PBST. Subsequently, counterstaining was performed using DAPI (4’,6-Diamidino-2-Phenylindole, Dihydrochloride, Thermo Fisher Scientific, Cat# D1306) (1μg/ml) and To-Pro-3 to visualize the nuclei of tissues. Samples were then washed three times and finally immersed in DABCO (1,4-diazabicyclo [2.2.2]octane, Sigma, Cat# D27802, 2.5% DABCO in 70% glycerol made in 1X PBS) until they were mounted on glass slides.

All antibodies were diluted in a blocking buffer. The following primary antibodies were used: mouse anti-γH2Av (1:1000, UNC93-5.2.1-s, DSHB)^36^, mouse anti-Hnt (1:100, 1G9c, DSHB), mouse anti-MMP1 catalytic domain (a cocktail of three antibodies at dilution 1:10, 3A6B4, 3B8D12, 5H7B11, DSHB), mouse anti-Draper (1:10, 5D14-s, DSHB)^90^, rabbit anti-cleaved Dcp-1 (1:100, 9578S, CST), rabbit anti-p-Akt (1:100, S473, CST), rabbit anti-phospho-ATM/ATR Substrate Motif (1;100, 6966S, CST), rabbit anti-Atg8a (1:100, Ab109364, Abcam), rabbit-pChk1 (1:100, Ab47318, Abcam), rabbit anti-histone H2AvD phosphoS137 (1:100, 600-401-914, Rockland) mouse anti-P1 (1:100, Istvan Ando)^64^ and rabbit anti-GFP(1:300, A11122, Invitrogen) rabbit anti-p-ASK1 (Thr845) (1:100, 3665S, CST). The secondary antibodies used for the immunohistochemistry are as follows: donkey anti-mouse AlexaFluor555 (A31570), goat anti-mouse AlexaFluor647 (A21050), donkey anti-rabbit AlexaFluor555 (A31572) and Goat anti-rabbit AlexaFluor647 (A32733) from Invitrogen and goat anti-mouse Cy3 (AB_2338680) from Jackson Scientific. All the secondary antibodies were used in 1:200 dilutions.

#### Dihydroethidium (DHE) staining for ROS

DHE staining Reactive Oxygen Species (ROS) was done as described in Owusu-Ansah and Banerjee, 2009^34^. Briefly, the lymph gland was isolated in Schneider’s *Drosophila* medium (Gibco, Cat# 21720024) at room temperature. The DHE (Dihydroethidium) dye (Invitrogen Molecular Probes, Cat# D11347) was prepared by reconstituting it in anhydrous DMSO (Sigma, Cat# D12345). The reconstituted DHE dye was dissolved in Schneider’s medium to achieve a final concentration of 1 μM/ml. Subsequently, the tissues were incubated in DHE dye for 5 min at room temperature, followed by three washes for 5 min each with Schneider’s medium. Finally, the tissues were mounted in DABCO, and images were acquired immediately.

#### Nick translation

The lymph glands were dissected in chilled 1X PBS, fixed in 4% paraformaldehyde for 30 min, and washed thrice for 10 min each with a wash buffer. Following these washes, tissues were rinsed with PBS supplemented with magnesium chloride (0.5mM) for 10 min each. The samples were transferred to PCR tubes and placed in a thermocycler at 37°C for 1 h. During this time, they were immersed in a reaction mixture consisting of 40 units/ml of *E. coli* DNA polymerase I (NEB, cat# M0209S), 50μM dATP, 50μM dGTP, 50μM dCTP, 35μM dTTP (Deoxynucleotide Set, 100mM, Sigma, Cat# DNTP100-KT), and 15μM DIG-11-dUTP (Digoxigenin-11-dUTP, alkali-labile, Sigma, Cat# 1157315291) in a 1X DNA polymerase reaction buffer. Following incubation, the samples were washed twice with wash buffer. They were then incubated for 2-hour incubation with a blocking solution at room temperature and subsequently incubated with anti-digoxigenin-Rhodamine (Anti-digoxigenin-Rhodamine, Fab fragments, Sigma, Cat# 11207750910) (0.5μg/mL) in the blocking solution for 2 hours at room temperature. After incubation with anti-digoxigenin-rhodamine, the tissues underwent additional washes with wash buffer, and finally, the samples were stained with DAPI and mounted in DABCO mounting medium^41^.

#### TUNEL staining

The lymph glands were isolated in cold PBS, fixed in 4% paraformaldehyde at room temperature for 30 minutes, and washed 3 times with 0.3% PBST. TUNEL (terminal deoxynucleotidyl transferase-mediated deoxyuridine triphosphate nick-end labeling) staining was performed using In Situ Cell Death Detection Kit, TMR Red (Sigma, cat# 12156792910) according to the manufacturer’s protocol^28^. Control eye disc (*GMR>UAS hid rpr*) as cell death positive control and another set of the same genotype incubated without enzyme as a negative control.

#### *Drosophila* larval staging

Synchronized 1^st^ instar larvae were collected just after hatching^28^. For synchronization, flies were allowed to lay embryos for 12 hours on egg-laying plates. After a 12-hour period of egg collection, these embryos were incubated at 25°C for 12 h. Following this incubation, hatched larvae were removed from the plate using a paintbrush, leaving behind unhatched embryos. The remaining unhatched embryos were incubated for 30 minutes at 25°C. The newly hatched larvae were carefully transferred to fresh vials of normal laboratory food and transferred to a 29°C incubator. Different staged larvae at 38 hours after larval hatching (ALH), 48h ALH, and 74h ALH were collected for γH2Av staining. Lymph glands were isolated, and immunostaining was performed as described in the immunostaining section.

#### LysoTracker staining

The lymph glands were dissected in PBS at room temperature and incubated in LysoTracker solution (LysoTracker Deep Red, Invitrogen, Cat# L12492) for 5 min. After incubation, tissues were washed with PBS and fixed in 4% PFA^80^. After fixation, the tissues were washed twice with PBS for 5 min each, stained with DAPI (1 mg/ml), and then washed with PBS. The tissues were mounted in DABCO, and imaging was performed under a confocal microscope.

#### Circulating blood cells counting

Third instar (L3) wandering larvae (*L-CasExpress L-Trace*) were bled in 20μl PBS on a clean coverslip, and hemocytes were allowed to adhere to the coverslip for 30 min. The PBS was carefully removed, and the cells were fixed with 4% PFA for 30 minutes. Following fixation, hemocytes were washed twice with PBS, stained with DAPI, and subjected to additional PBS washes^38^. The prepared samples were then mounted on clean slides. Using a Zeiss LSM-900 confocal microscope with 10X and 20X objectives, three random images were captured for each larval bleeding sample, encompassing GFP and DAPI channels. The number of DAPI- and GFP-positive hemocytes from each image was quantified manually using ImageJ, and the percentage of GFP-positive hemocytes was obtained.

#### Circulating hemocyte immunostaining

Third instar (L3) wandering larvae were bled in 20μl PBS on a coverslip, and hemocytes were allowed to adhere to the coverslip for 30 min. The PBS was removed, and the cells were fixed with 4% PFA for 30 min. After fixation, immunostaining was performed similarly to lymph gland immunostaining as described earlier^38^.

#### Phagocytic assay of circulating hemocytes

RFP-expressing *E. coli* (Addgene Cat# 17827) bacterial culture obtained from overnight culture in LB broth (Luria Bertani broth, HIMEDIA, Cat# M1245) supplemented with 0.2 % L-Arabinose (Sigma, Cat# A81906) and 100μg/ml Ampicillin (Sigma, Cat# A5354), was taken in a clean microcentrifuge tube. Bacteria were precipitated using centrifugation, and precipitated bacteria were washed with PBS. After washing, the bacteria precipitate was suspended in 100μl of autoclaved PBS. 1 μl of this suspension was used in each experiment. Phagocytosis assay was conducted using circulatory hemocytes isolated from wandering third-instar larvae^92^. These hemocytes were collected by bleeding the larvae onto a coverslip, where they came into contact with RFP-expressing *E. coli* suspended in autoclaved PBS. After a 10-minute incubation in a humid chamber, the solution was removed, and the hemocytes were fixed using a 4% PFA fixative solution for 30 minutes. Following fixation, cells were washed with PBS two times, 10 minutes each, and subsequently, they were stained with DAPI (1μg/ml) for 30 minutes, washed with PBS twice, and mounted on a clean slide. For each larval bleeding sample, three random images were captured using a fluorescent microscope (Nikon E800) with a 20X objective lens for RFP, GFP, and DAPI channels. The number of hemocytes positive with bacteria and without bacteria was quantified by ImageJ manually, and phagocytic efficiency was calculated.

#### Embryo live imaging

*Drosophila* embryos at the desired developmental stage were collected from overnight eggs laying on apple juice plates. Subsequently, these embryos underwent a dechorionation process involving a 5-minute treatment with 4% bleach, followed by two rinses with 1X PBS. The dechorionated embryos were carefully positioned on a slide coated with halocarbon oil (Sigma-Aldrich Cat# H8898). The embryos were immersed in oil, and a cover glass was placed over them. Time-lapse imaging was performed using a confocal microscope, capturing images at 30-second intervals for both the green and red channels^97^. Different zoom settings were applied during the imaging process to obtain various magnification levels or fields of view as needed.

#### Microscopy and image processing

All samples were imaged in a Zeiss LSM-900 confocal microscope using Zen blue software under a 20X objective with a zoom of 1.0 and a 40X objective with a zoom of 0.5 and used a 2.0 optical section interval in all images otherwise specified in the figure legend. All images were processed using ImageJ software (NIH, USA) (available at imagej.nih.gov/ij), and Microsoft PowerPoint was used to make the figure panel. Images of LGs are a maximum intensity projection of the stack of the middle third of the samples; it allows for visibility of the inside of the LG, which can be covered by the cortical zone region in a maximum intensity projection of the entire LG, specified in the figure legend.

#### Quantification of lymph gland phenotypes

All quantification was done using ImageJ software (NIH, USA). The number of γH2Av, Hnt, and Dcp-1 positive cells and colocalization of γH2Av with Dcp-1, γH2Av with *GC3Ai*, Hnt with *L-CasExpress L-Trace* was counted manually for both lobes of the primary lymph gland, and analyzed separately. For volume measurement of multichannel images, first, all channels of images were separated, and one specific threshold was chosen that fit best for the actual staining and kept constant throughout the measurement. The thresholding procedure is used in image processing to select pixels of interest based on the intensity of the pixel values. After that, the “Measure stack” plugin^59^ was used to find the fluorescent area (DAPI, GFP, Draper, and P1) of each optical section. Then, the fluorescent area in each optical section was added and multiplied with stack interval (2µm) to find out the volume. For better representation, the primary lobe of the lymph gland has been represented and outlined in white dashed lines.

#### RNA isolation, quantitative reverse transcription PCR analysis

Total RNA was isolated from thirty wandering 3^rd^ instar larval fat body using Trizol reagent following the manufacturer’s recommended protocol (Sigma-Aldrich, Cat#T9424). The RNA pellets were resuspended in 15μl of DEPC-MQ water, and after the pellets were dissolved, their quantitative estimation was done using spectrophotometric analysis. Subsequently, 1μg of each RNA sample was incubated with 1U of RNase-free DNase I (Thermo Fisher Scientific, Cat# 89836) for 30 minutes at 37°C to eliminate residual DNA. Following the standard cDNA preparation protocol, the first-strand cDNA was synthesized from these incubated samples. The prepared cDNA was subjected to a real-time PCR machine using forward and reverse primer pairs of the target genes. Real-time PCR was done by using 5 μl of qPCR master mix (SYBR Green, Genetix, Cat# PKG025-A), 2 picomol/μl of each primer per reaction in 10μl of final volume in ABI 7500 real-time PCR machine (Applied Biosystems). The relative fold change in mRNA expression for different genes was calculated using the comparative C_T_ method to assess changes in gene expression. Data normalization was done using *Rp49* as an internal control. For each gene, three independent biological replicates were used. The following primers are used for this study:

*Drep1* (Forward) 5’-GCGACCACCTGCCCACCATCCAC-3’
*Drep1* (Reverse) 5’-CTCCGTTTCCAGTTCCTGCCTCG-3’
*Drep4* (Forward) 5’- CAGCCAACAGGCGAAGTTCTCAAGG-3’
*Drep4* (Reverse) 5’- CGGGCCAGACACTATGAACAGCGTC-3’
*Rp49* (Forward) 5’- TTGAGAACGCAGGCGACCGT-3’
*Rp49* (Reverse) 5’- CGTCTCCTCCAAGAAGCGCAAG-3’

#### Statistical analysis

All experiments have been repeated at least three times, from which one representative image is shown. The quantifications shown are for all the sets analyzed. All the statistical tests for the respective experiments were carried out using Microsoft Excel 2019 and GraphPad Prism 9. All the p-values represent unpaired two-tailed Student’s t-tests to determine statistical significance. The significance level is indicated by an * for p ≤ 0.05, ** for p ≤ 0.01, *** for p ≤ 0.001, **** for p ≤ 0.0001, and by ns for not significant, p > 0.05.

## Supplementary figure legends

**Figure S1.**
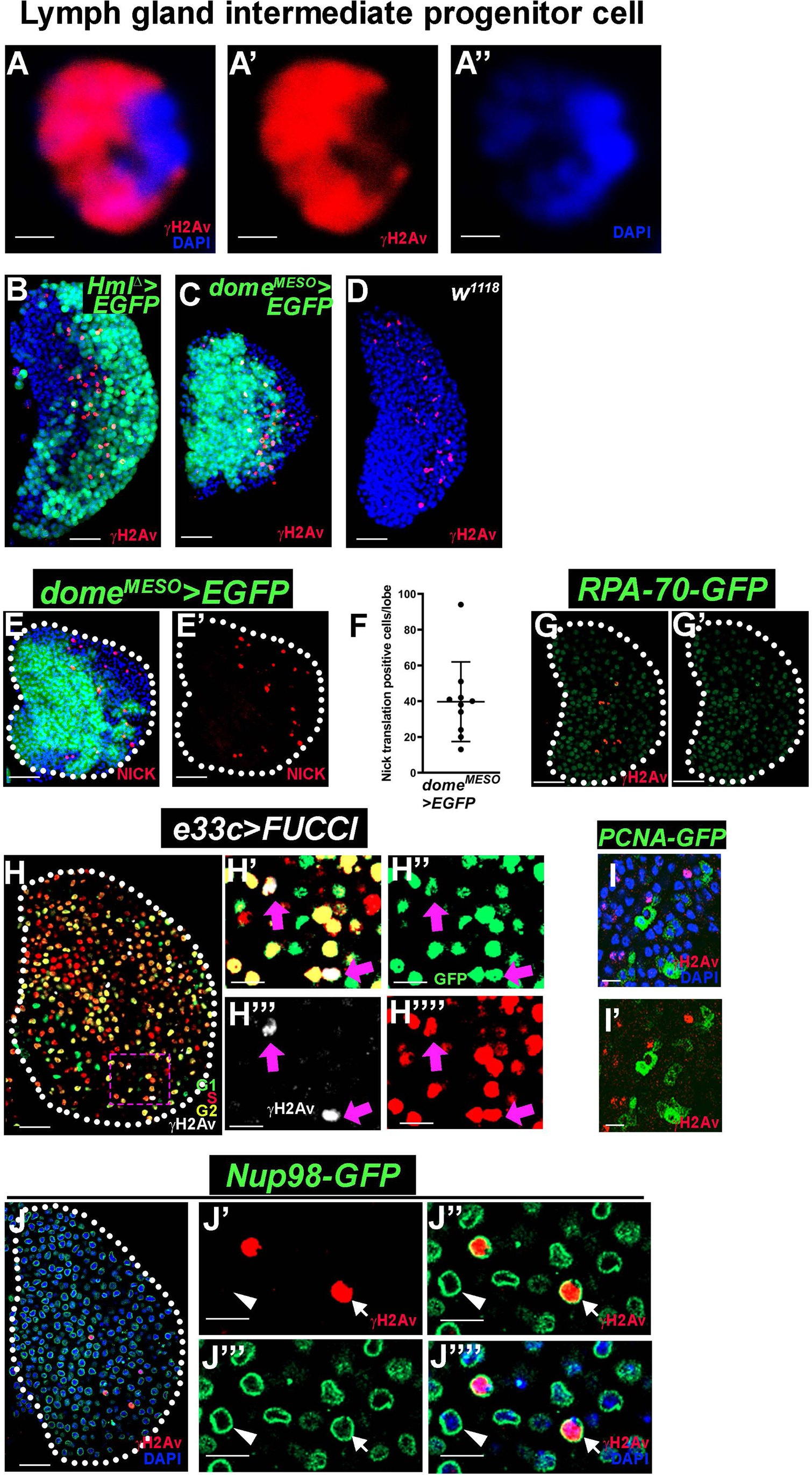
DNA damage occurs during the differentiation of lymph gland. **(A-A’’)** The γH2Av staining (red) covers the entire nuclear region, except the DAPI (blue)-bright heterochromatin region in the lymph gland intermediate progenitor. **(B-D)** *Hml^Δ^-Gal4, UAS-2xEGFP/+* (green) (B), *dome^MESO^-Gal4, UAS-2xEGFP*/+ (green) (C) and *w^1118^* (D) genetic backgrounds show a similar number of γH2Av positive cells (red) in the differentiating zone. **(E-E’)** Nick translation (red) shows in differentiating cells (low GFP) of control lymph gland indicating DNA strand breaks *dome^MESO^-Gal4, UAS-2xEGFP/+* (green) (n=10). **(F)** Quantification of nick translation positive cell number in control (*dome^MESO^-Gal4, UAS-2xEGFP/+*) (green) represented in (E-E’). **(G-G’)** γH2Av staining (red) colocalized with *RPA70-GFP* (green) in lymph gland. **(H-H’’’’)** Expression of *e33c-Gal4, UAS-FUCCI* shows the cell cycle status of lymph gland G2 (yellow), G1 (green), and S (red) phases (H) and colocalization of G2 cells with γH2Av (grey). The insets show the colocalization of γH2Av -positive cells with GFP and RFP, indicated by the arrows in the G2 phase (H’-H””). **(I-I’)** γH2Av staining (red) in *PCNA-GFP* (green) cells as S phase marker do not colocalize in the lymph gland, nuclei stained with DAPI (blue). **(J-J’’’’)** γH2Av staining (red) in *Nup98-GFP* (green) with DAPI (blue) (J) and high magnification image of γH2Av in *Nup-98-GFP* background only γH2Av shows intact nuclear membrane (J’), γH2Av with *Nup98-GFP* (J’’), only *Nup98-GFP* (J’’’) and γH2Av, *Nup98-GFP* with DAPI (J’’’’). Arrow marks the γH2Av positive nuclei and intact nuclear pore complexes compared with γH2Av negative (marked arrowhead) nuclei in the lymph gland. All images are from wandering third instar lymph gland. Scale bars: 25µm in all images except 10µm in (H’-I’ and J’-J’’’’), and 1µm in (A-A’), with single optical sections except image (B-E’), which are maximum intensity projections of the middle third optical sections. Nuclei stained with DAPI (blue). Lymph glands boundary demarcated by white dotted line for clarity. Error bars, mean ± SD. All images are representative of 3 or more independent biological experiments.

**Figure S2.**
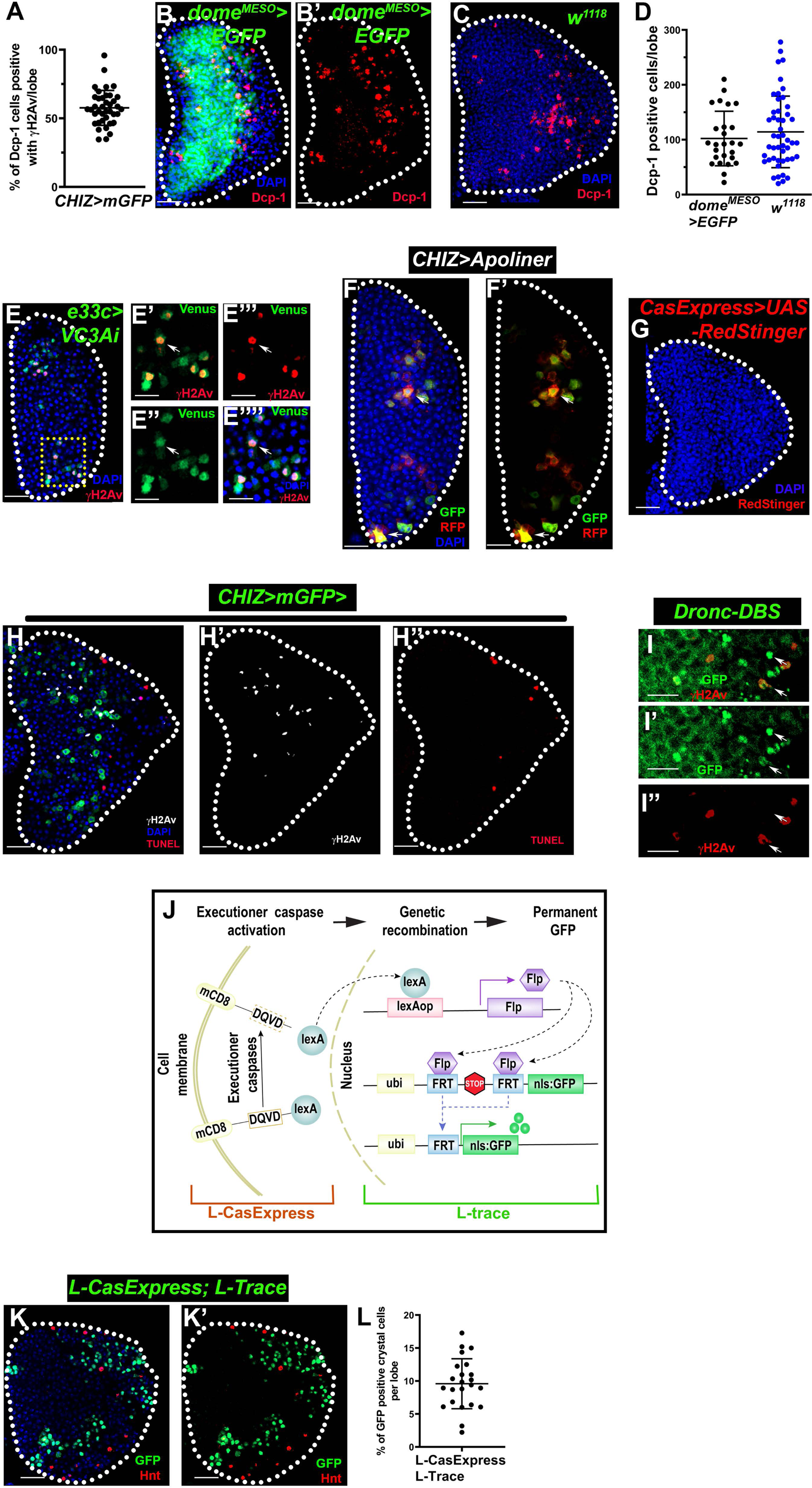
DNA breaks and active caspase in differentiating progenitor blood cells. **(A)** Quantification of the percentage of Dcp-1 positive cells colocalized with γH2Av positive cells in the control lymph gland per lobe [*CHIZ>mGFP/+* (n=39)]. **(B-C)** Dcp-1 staining (red) is shown in *dome^MESO^-Gal4, UAS-2xEGFP/+* progenitors mark green (B-B’) and *w^1118^* (C) genetic background of lymph gland in the intermediate zone. **(D)** Quantification of Dcp-1 positive cell number in *dome^MESO^-Gal4, UAS-2xEGFP/+* (n=26), and *w^1118^* background (n=48) per lymph gland lobe show a similar pattern. **(E-E’’’’)** Fluorescent reporter (green) of executioner caspase (*e33c-Gal4, UAS-VC3Ai*) are shown with γH2Av staining (red) (E). Insets show venus (green) colocalization with γH2Av (arrow). **(F-F’)** Expression of *CHIZ-Gal4, UAS-Apoliner* show membrane RFP and membrane GFP as well as nuclear GFP (marks caspase active cell), arrow indicates caspase active cell. **(G)** Expression of *CasExpress^Mutant^-Gal4, UAS-RedStinger* does not show executioner activity in the lymph gland, which is used as a control for *CasExpress-Gal4* (Figure 2H). **(H-H’’)** γH2Av-positive cells (grey) in the lymph gland are not high-intensity TUNEL-positive cells (red). **(I-I”)** High-magnification image of γH2Av (red) staining in *Drice-Based-Sensor-GFP* (*DBS-GFP*; green) (I), only γH2Av (I’) and only GFP (I’’) shows γH2Av colocalization with low DBS-GFP cells. **(J)** A model showing the mechanistic principle underlying how the *L-CasExpress L-Trace* works (adapted from Sun G *et al.* 2021). **(K-K’)** Crystal cell marker Hnt staining (red) in the *L-CasExpress L-Trace* (*lex-Aop-Flp::Ubi-FRT-STOP-FRT-GFP/+; L-caspase/+*) show significantly less colocalization with lineage trace GFP (K) and only Hnt (K’) **(L)** Quantification of the percentage of crystal cells in *L-CasExpress L-Trace* (*lex-Aop-Flp::Ubi-FRT-STOP-FRT-GFP/+; L-caspase/+*) colocalized with lineage trace GFP(green) and DAPI (blue) (n=23). All images are from wandering third instar lymph gland. Scale bars: 25µm in all images except 10µm in (E-E’’’’ and I-I’’). All are maximum intensity projections of the middle third optical sections except images (E-F’’, H-I’’ and K-K’), which are single optical sections. Nuclei stained with DAPI (blue). Lymph glands boundary demarcated by white dotted line for clarity. Error bars, mean ± SD. All images are representative of 3 or more independent biological experiments.

**Figure S3.**
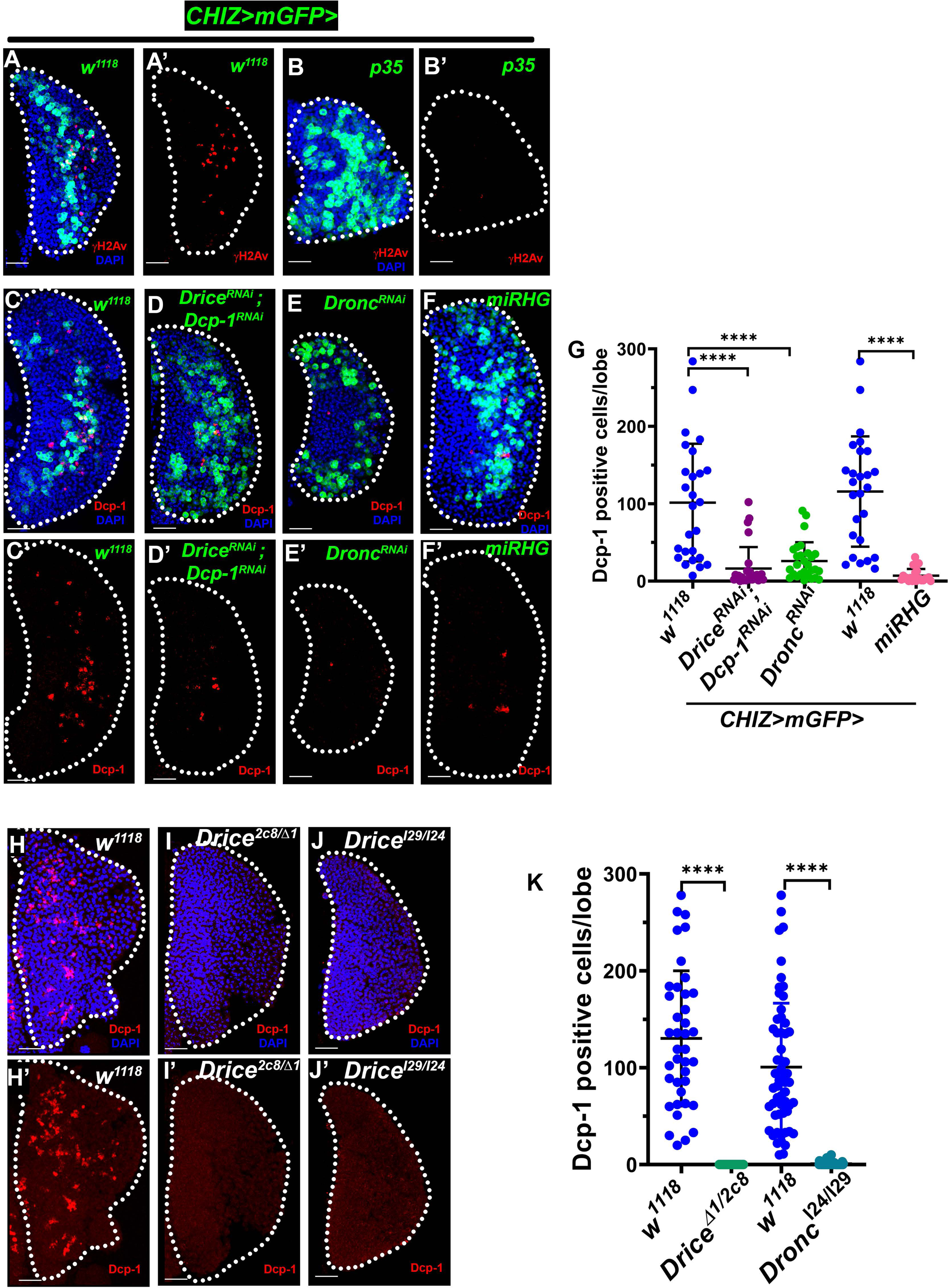
Caspase-mediated DNA breaks are essential for macrophage differentiation. **(A-B’)** γH2Av staining (red) in *CHIZ>mGFP* driven *UAS-p35* background shows very less γH2Av positive cell as compared to control (A-A’). **(C-F’)** One representative image of control, *CHIZ>mGFP/+* (C-C’), *CHIZ>mGFP* driven *RNAi, UAS-Drice^RNAi^; UAS-Dcp-1^RNAi^* (D-D’)*, UAS-Dronc^RNAi^* (E-E’)*, UAS-miRHG* (F-F’) with Dcp-1 (red) staining show less Dcp-1 positive cells. **(G)** Quantification of Dcp-1 positive cell number in control, *CHIZ>mGFP/+* (n=26) with UAS-*Drice^RNAi^; UAS-Dcp-1^RNAi^* (n=28) and *UAS-Dronc^RNAi^* (n=28); control, *CHIZ>mGFP/+* n=26) with *UAS-miRHG* (n=23) shows significantly reduced Dcp-1 positive cells. Here, control sets differ for each *RNAi* because one set of experiments with a particular gene RNAi was done separately. **(H-J’)** Dcp-1 staining (red) in control *w^1118^* (H-H’), but mutants *Drice^Δ1^/Drice^2c8^* (I-I’) and *Dronc^I29^/Dronc^I24^* (J-J’) and mutants do not show Dcp-1 positive cells. **(J)** Quantification of Dcp-1 positive cell number in control *w^1118^*(n=39) with *Drice^Δ1^/Drice^2c8^* (n=41) and control *w^1118^* (n=59) with *Dronc^I29^/Dronc^I24^*(n=58), mutants show no Dcp-1 positive cells. Here, control sets differ for each *RNAi* because one set of experiments with a particular RNAi was done separately. All images show a 25µm scale bar, with maximum intensity projections of the middle third optical section of the wandering third instar larval lymph gland lobe. Nuclei stained with DAPI (blue). Lymph glands boundary demarcated by white dotted line for clarity. ****P < 0.0001 Error bars, mean ± SD. All images are representative of 3 or more independent biological experiments.

**Figure S4.**
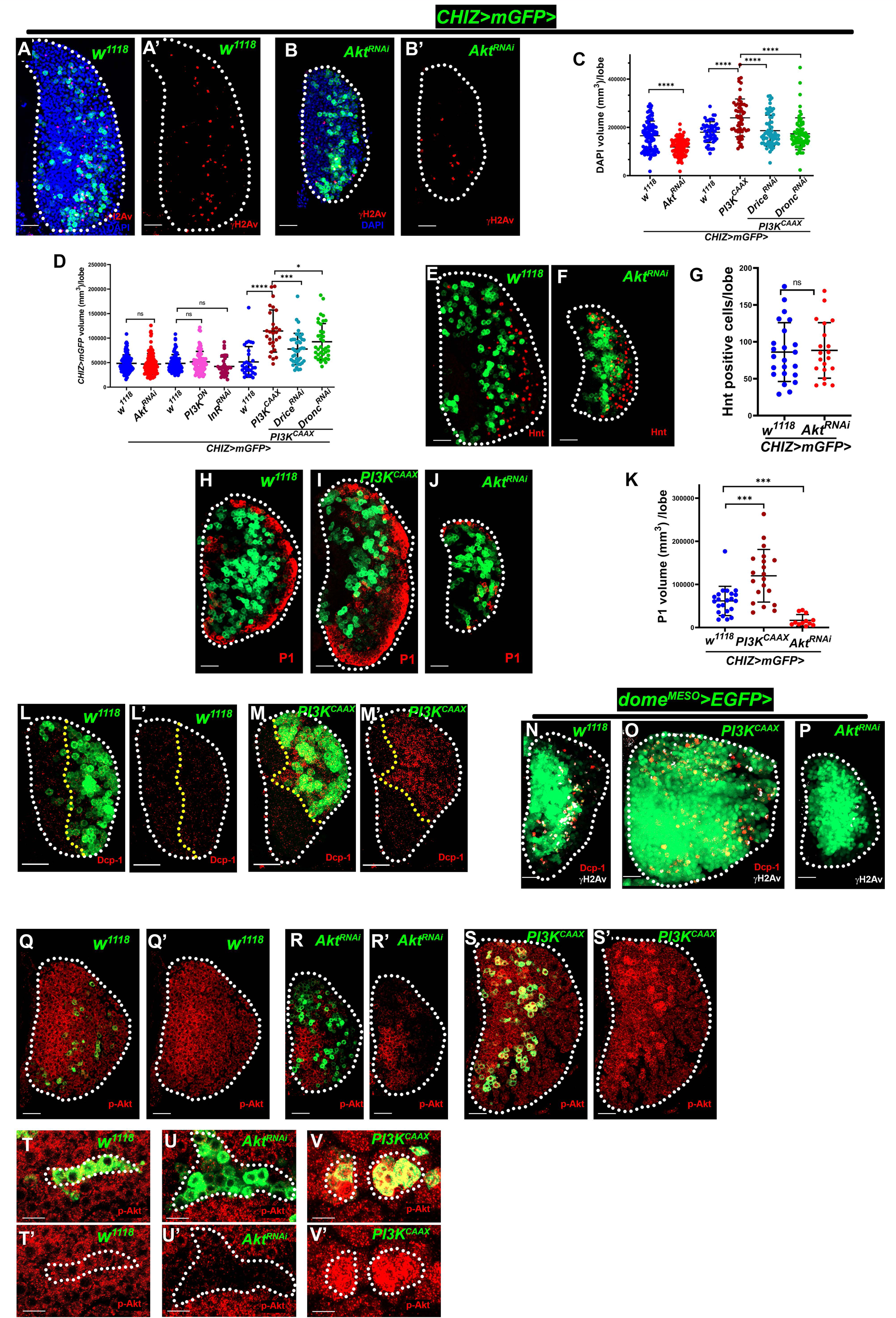
InR-PI3K-Akt signaling regulates caspase activity and DNA breaks in macrophage differentiation. **(A-B’)** *CHIZ>mGFP* (green) driven *Akt^RNAi^* (BL 31701) (B-B’) has less γH2Av positive cells (red) as compared to control (A-A’). **(C)** Quantification of lymph gland size by measuring the DAPI volume in control, *CHIZ>mGFP/+* (96) and significantly reduced in *Akt^RNAi^* (n=102), control *CHIZ>mGFP/+* (n=50) significantly high in *PI3K^CAAX^* (n=58), and rescued in *PI3K^CAAX^; Drice^RNAi^* (n=63) and *PI3K^CAAX^*; *Dronc^RNAi^* (n=69). Here, control sets are different for groups of experimental sets because experimental sets are done on different days. **(D)** Quantification of the GFP positive cells volume in control, *CHIZ>mGFP/+* (122) and *CHIZ>mGFP* driven *Akt^RNAi^* (n=119), control *CHIZ>mGFP/+* (n=83) and *PI3K^DN^* (n=78), *InR^RNAi^*(n=34) show no change in GFP+ cells volume. However, *CHIZ>mGFP* driven *PI3K^CAAX^* (n=28) show high GFP+ cells’ volume and partially rescued in *PI3K^CAAX^; Drice^RNAi^*(n=41) and *PI3K^CAAX^*; *Dronc^RNAi^* (n=38) compared to control *CHIZ>mGFP/+* (n=30) group. Here, control sets are different for groups of experimental sets because experimental sets are done on different days. **(E-F)** *CHIZ>mGFP* driven *Akt^RNAi^* (F) do not show the difference in crystal cells (Hnt staining in red) number compared to in control, *CHIZ>mGFP/+* (E). **(G)** Quantification of the Hnt positive cell number in control, *CHIZ>mGFP/+* (n=23), and *CHIZ>mGFP* driven *Akt^RNAi^* (n=20) do not show differences in crystal cell numbers. **(H-J)** *CHIZ>mGFP* driven *PI3K^CAAX^*(I) shows high P1 (red), a macrophage marker compared to control, *CHIZ>mGFP*/+ (H), but *CHIZ>mGFP* driven *Akt^RNAi^* (J) shows less P1. **(K)** Quantification of the P1 volume in control, *CHIZ>mGFP/+* (n=22), and *CHIZ>mGFP* driven *PI3K^CAAX^* (n=20) show a significantly high number of macrophages (P1 positive cells), *CHIZ>mGFP* driven *Akt^RNAi^* (n=12) show significantly less P1. **(L-M’)** Dcp-1 staining (red) in control, *CHIZ>mGFP/+* (L-L’) and *CHIZ>mGFP* driven *PI3K^CAAX^* (M-M’) show high Dcp-1 staining in early third instar larvae (48 hours after hatching at 29°C) in CHIZ+ cells. **(N-P)** γH2Av (grey) and Dcp-1 (red) co-staining in control *dome^MESO^-Gal4, UAS-2xEGFP/+* (N) and *dome^MESO^-Gal4, UAS-2xEGFP/+* driven *PI3K^CAAX^* (O) where high number of γH2Av and Dcp-1 positive cells found only in low GFP cells, whereas *dome^MESO^-Gal4, UAS-2xEGFP/+* driven *Akt^RNAi^* (P) show less γH2Av positive cells. **(Q-S’)** Lymph gland exhibit active Akt signaling shown by p-Akt staining (red) in control, *CHIZ>mGFP/+* (Q-Q’) (green), but *CHIZ>mGFP* driven *Akt^RNAi^* (R-R’), show less p-Akt staining, and *PI3K^CAAX^*(S-S’) show high p-Akt staining. **(T-V’)** p-AKT staining (red) in control, *CHIZ>mGFP/+* (T-T’), and *CHIZ>mGFP* driven *Akt^RNAi^* (U-U’) shows less staining in CHIZ^+^ cells, and *PI3K^CAAX^* (V-V’) shows high staining in CHIZ^+^ cells. All images are from wandering third instar lymph gland lobes except image L-M’, the early third instar lymph gland lobe. Scale bars: 25µm in all images except 10µm in (T-V’) with maximum intensity projections of the middle third optical sections except image Q-V’, which are single optical sections. Nuclei stained with DAPI (blue). Lymph glands boundary demarcated by white dotted line for clarity. *P < 0.05 ***P < 0.001, ****P < 0.0001, ns-not significant Error bars, mean ± SD. All images are representative of 3 or more independent biological experiments.

**Figure S5.**
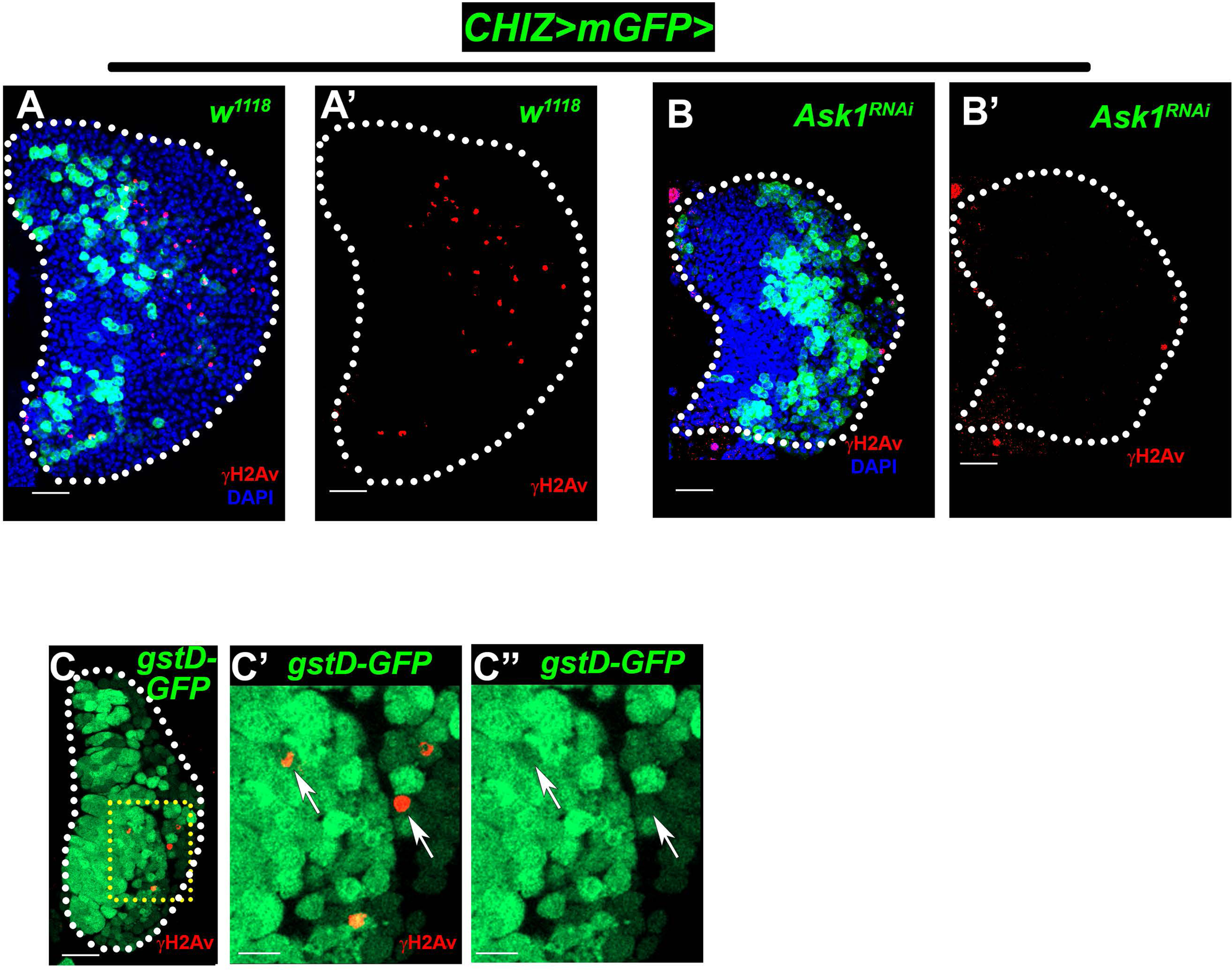
Caspase activity regulated by attenuated Ask1. **(A-B’)** *CHIZ>mGFP* driven *Ask1^RNAi^* (BL32646) (B-B’) shows reduce γH2Av positive cells (red) as compared to control *CHIZ>mGFP* (A-A’). **(B-B’’)** γH2Av (red) positive cells in the lymph gland display low *gstD-GFP/+* (green) (B), and high magnification images (B’-B’’) show γH2Av positive cells in low gstD-GFP cells indicated by arrow. All images are from the wandering third instar lymph gland lobe show a 25µm scale bar except image (C’-C’’), where scale bar is 10µm and maximum-intensity projections of the middle third optical sections except images C-C’’, which are single optical sections. Nuclei stained with DAPI (blue). Lymph glands boundary demarcated by white dotted line for clarity. All images are representative of 3 or more independent biological experiments.

**Figure S6.**
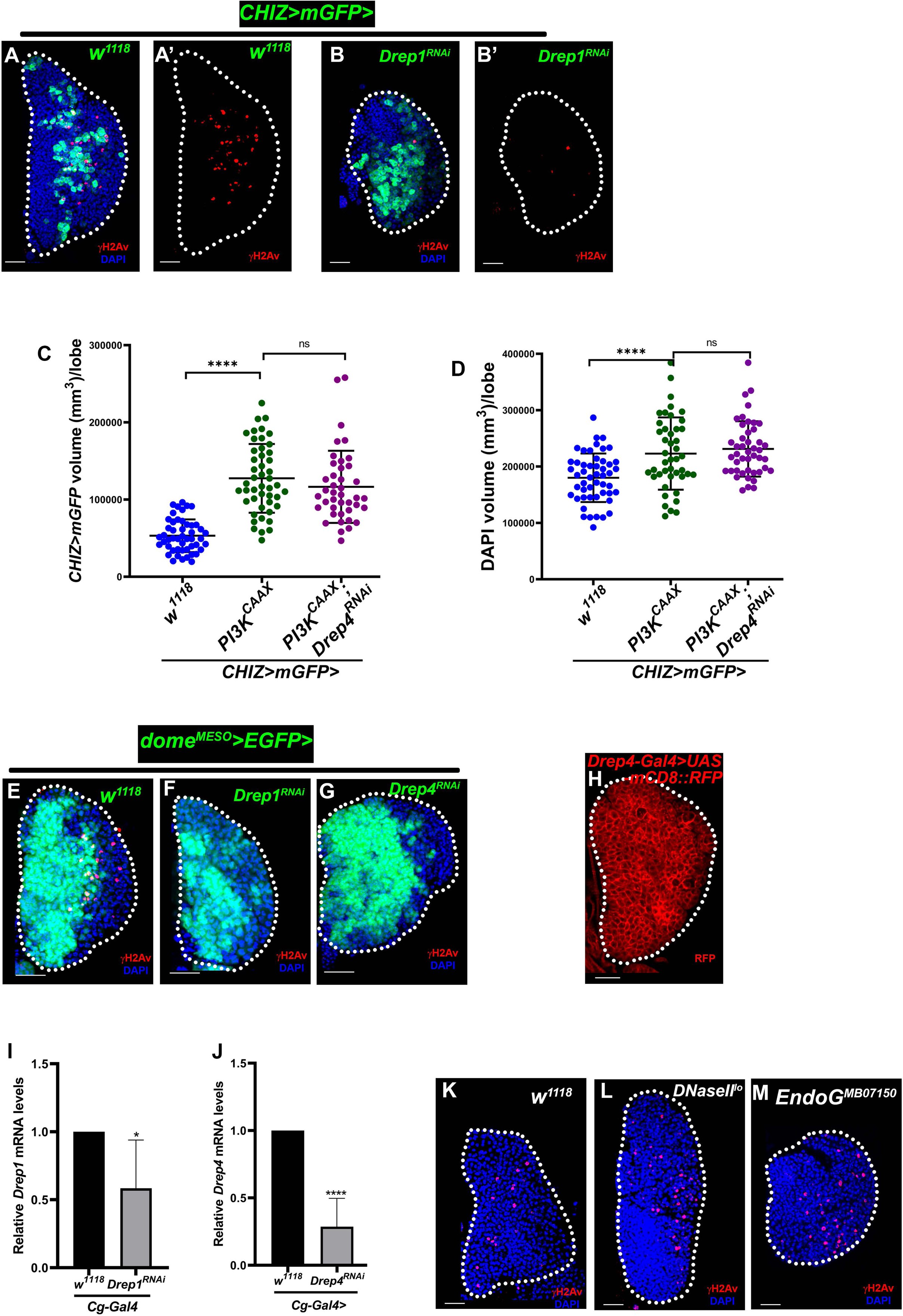
Caspase Activated DNase (CAD) induced DNA breaks required for macrophage differentiation. **(A-B’)** Depletion of *Drosophila* ICAD [(*CHIZ>mGFP; UAS*-*Drep1^RNAi^* (BL65944) (B-B’) in the intermediate progenitors cause less γH2Av positive cells (red) GFP cell marks intermediate progenitors (green) compared to control *CHIZ>mGFP/+* (A-A’). **(C)** Quantification of GFP volume in control*, CHIZ>mGFP/+* (n=49) and *CHIZ>mGFP* driven *UAS*-*PI3K^CAAX^* (n=47) have significantly high GFP volume*, UAS*-*PI3K^CAAX^; UAS-Drep4^RNAi^* (n=41) rescue high GFP volume. **(D)** Quantification of DAPI volume in control, *CHIZ>mGFP/+* (n=50) and *CHIZ>mGFP* driven *UAS*-*PI3K^CAAX^* (n=44) have significantly high GFP volume*, UAS*-*PI3K^CAAX^; UAS-Drep4^RNAi^* (n=45) rescue high GFP volume. **(E-G)** Depletion of *Drosophila* ICAD (*dome^MESO^-Gal4, UAS-2xEGFP; UAS*-*Drep1^RNAi^ VDRC8357*) (F) and CAD (*dome^MESO^-Gal4, UAS-2xEGFP*; *UAS-Drep4^RNAi^*) (G) in the progenitors lead to significantly less γH2Av positive cells (red) in the lymph glands compared to control, *dome^MESO^-Gal4, UAS-2xEGFP*/+ (E). GFP cell marks progenitors (green). **(H)** *Drosophila* CAD (Drep4) expressed (*Drep4-Gal4, UAS-mCD8::RFP)* in wandering third instar larval lymph gland. **(I-J)** Quantitate RT-PCR show that Drep1 (ICAD) (I) and Drep4 (CAD) (J) expression in larval fat body and RNAi lines targeting Drep1 and Drep4 by using *Cg-Gal4* reduces both transcript levels. **(K-M)** γH2Av staining (red) in control, both mutants *DNaseII^lo^* (L) *and EndoG^MB07150^* (M) have γH2Av positive cells similar to control *w^1118^* (K) in the lymph gland. All images are from wandering third instar lymph gland lobes show a 25µm scale bar and maximum intensity projections of the middle third optical section except image H, which is a single optical section. Nuclei stained with DAPI (blue). Lymph glands boundary demarcated by white dotted line for clarity. ***P < 0.001, ****P < 0.0001, ns-not significant Error bars, mean ± SD. All images are representative of 3 or more independent biological experiments.

**Figure S7.**
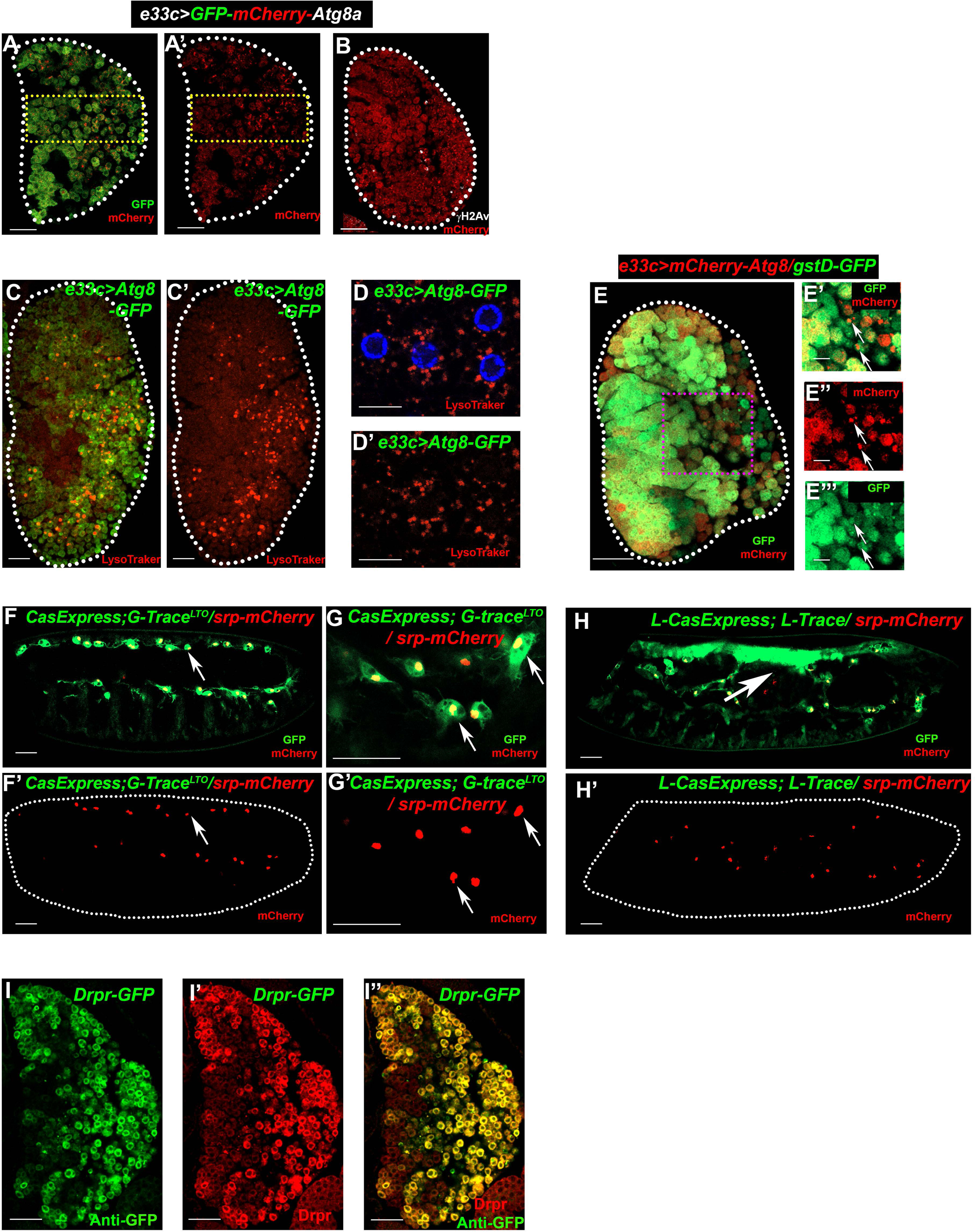
Developmental caspase-activated DNA breakage and autophagy regulate macrophage differentiation. **(A-B)** *e33c-Gal4, UAS-GFP-mCherry-Atg8a* expression shows *Atg8a-mCherry* puncta in lymph gland (A-A’), colocalized in the γH2Av positive cells (B), also see Figure 7. **(C-D’)** *e33c-Gal4, UAS Atg8a-GFP* expression shows LysoTracker staining (red) with GFP (C) and only LysoTracker (C’) in the intermediate to differentiated zone. Also, in high magnification image of fat body as a control (D-D’). **(E-E’’’)** Expression of *e33c-Gal4 UAS*-*mCherry*-*Atg8a* in with *gstD-GFP* (E). Insets show the high *Atg8a-mCherry* puncta in low ROS (green) cells (E’-E’’’) marked by arrow. **(F-G’)** Embryonic macrophages (stage 13) marked by *srp-mCherry* shows experience of caspase lineage activity (*CasExpress; G-Trace^LTO^*) (F-F’) also shows in high magnification (G-G’), marked by arrows. **(H-H’)** Embryonic macrophages (stage 13) marked by *srp-mCherry* experience caspase lineage (*L-CasExpress L-Trace*) GFP (green) dorsal closure (marked by arrow) and macrophages. **(I-I’’)** *Draper-GFP* expression using anti-GFP (green) (G) and staining of anti-Draper (red) antibody (G’) and merge image of both (G’’). All images are from wandering third instar lymph gland lobes, except F-H’. All images show a 25µm scale bar except image (D-D’ and G-G’) where the scale bar is 10µm and image E’-E’’ have 5µm scale bar. All images are single optical sections. Nuclei stained with DAPI (blue). Lymph glands boundary demarcated by white dotted line for clarity. All images are representative of 3 or more independent biological experiments. Also see Movies S1, S2 and S3.

## Supplementary legends for movies

**Movie S1.** Executioner caspase activates during the embryonic macrophage development, related to **Figure S7**. Time-lapse (30-second interval) movie from the ventral region of the late embryo (stage 13). Embryonic macrophage nuclei labeled in mCherry (*srp-mCherry*) and the cytoplasm in green (GFP) expressed in the active caspase Lineage Trace Only (LTO) cells (*CasExpress-Gal4; G-Trace^LTO^*). All embryonic macrophages (*srp-mCherry* positive cells) experience caspase activity during their development, resulting in GFP expression in their lineage cells.

**Movie S2.** Active macrophages experience the executioner caspases activity, **Figure S7**. High magnification time-lapse (20-second interval) movie from the posterior ventral area of the late embryo (stage 13). Embryonic macrophage nuclei labeled in mCherry (*srp-mCherry*) and the cytoplasm in green (GFP) expressed in the active caspase lineage cells (*CasExpress-Gal4; G-Trace^LTO^*). All embryonic macrophages (*srp-mCherry* positive cells) experience caspase activity during their development, resulting in GFP expression in their lineage cells. These embryonic macrophages show active pseudopodia movement and contain vacuoles, suggesting their proper development as phagocytic macrophages.

**Movie S3.** Activation of caspase during the development of *Drosophil*a embryonic macrophages, related to Figure 7. Time-lapse (30-second interval) movie from the lateral view of the late embryo (stage 13). Embryonic macrophage nuclei labeled in mCherry (*srp-mCherry*) and the cytoplasm in green (GFP) expressed in the active caspase lineage cells (*L-CasExpress L-Trace*). Embryonic macrophages (*srp-mCherry* positive cells) experience caspase activity during their development, resulting in GFP expression in their lineage cells. During the *Drosophila* embryonic development, caspases are activated in many other cells, especially in the dorsal closure region, where the intense lineage GFP is expressed (marked by arrow).

## References

1. Lazarov, T., Juarez-Carreno, S., Cox, N., and Geissmann, F. (2023). Physiology and diseases of tissue-resident macrophages. Nature 618, 698–707. 10.1038/s41586-023-06002-x.

2. Park, M.D., Silvin, A., Ginhoux, F., and Merad, M. (2022). Macrophages in health and disease. Cell 185, 4259–4279. 10.1016/j.cell.2022.10.007.

3. Mantovani, A., Allavena, P., Marchesi, F., and Garlanda, C. (2022). Macrophages as tools and targets in cancer therapy. Nat Rev Drug Discov 21, 799–820. 10.1038/s41573-022-00520-5.

4. Cox, N., Pokrovskii, M., Vicario, R., and Geissmann, F. (2021). Origins, Biology, and Diseases of Tissue Macrophages. Annu Rev Immunol 39, 313–344. 10.1146/annurev-immunol-093019-111748.

5. Gosselin, D., Link, V.M., Romanoski, C.E., Fonseca, G.J., Eichenfield, D.Z., Spann, N.J., Stender, J.D., Chun, H.B., Garner, H., Geissmann, F., and Glass, C.K. (2014). Environment drives selection and function of enhancers controlling tissue-specific macrophage identities. Cell 159, 1327–1340. 10.1016/j.cell.2014.11.023.

6. Lavin, Y., Winter, D., Blecher-Gonen, R., David, E., Keren-Shaul, H., Merad, M., Jung, S., and Amit, I. (2014). Tissue-resident macrophage enhancer landscapes are shaped by the local microenvironment. Cell 159, 1312–1326. 10.1016/j.cell.2014.11.018.

7. Wang, Z., Wu, Z., Wang, H., Feng, R., Wang, G., Li, M., Wang, S.Y., Chen, X., Su, Y., Wang, J., et al. (2023). An immune cell atlas reveals the dynamics of human macrophage specification during prenatal development. Cell. 10.1016/j.cell.2023.08.019.

8. Coillard, A., and Segura, E. (2019). In vivo Differentiation of Human Monocytes. Front Immunol 10, 1907. 10.3389/fimmu.2019.01907.

9. Alvarez-Errico, D., Vento-Tormo, R., Sieweke, M., and Ballestar, E. (2015). Epigenetic control of myeloid cell differentiation, identity and function. Nat Rev Immunol 15, 7–17. 10.1038/nri3777.

10. Tiwari, S.K., Toshniwal, A.G., Mandal, S., and Mandal, L. (2020). Fatty acid beta-oxidation is required for the differentiation of larval hematopoietic progenitors in Drosophila. Elife 9. 10.7554/eLife.53247.

11. Wculek, S.K., Dunphy, G., Heras-Murillo, I., Mastrangelo, A., and Sancho, D. (2022). Metabolism of tissue macrophages in homeostasis and pathology. Cell Mol Immunol 19, 384–408. 10.1038/s41423-021-00791-9.

12. Clarke, A.J., and Simon, A.K. (2019). Autophagy in the renewal, differentiation and homeostasis of immune cells. Nat Rev Immunol 19, 170–183. 10.1038/s41577-018-0095-2.

13. Solier, S., Fontenay, M., Vainchenker, W., Droin, N., and Solary, E. (2017). Non-apoptotic functions of caspases in myeloid cell differentiation. Cell Death Differ 24, 1337–1347. 10.1038/cdd.2017.19.

14. Hoeffel, G., Chen, J., Lavin, Y., Low, D., Almeida, F.F., See, P., Beaudin, A.E., Lum, J., Low, I., Forsberg, E.C., et al. (2015). C-Myb(+) erythro-myeloid progenitor-derived fetal monocytes give rise to adult tissue-resident macrophages. Immunity 42, 665–678. 10.1016/j.immuni.2015.03.011.

15. Green, D.R. (2019). The Coming Decade of Cell Death Research: Five Riddles. Cell 177, 1094–1107. 10.1016/j.cell.2019.04.024.

16. Burgon, P.G., and Megeney, L.A. (2018). Caspase signaling, a conserved inductive cue for metazoan cell differentiation. Semin Cell Dev Biol 82, 96–104. 10.1016/j.semcdb.2017.11.009.

17. Fuchs, Y., and Steller, H. (2011). Programmed cell death in animal development and disease. Cell 147, 742–758. 10.1016/j.cell.2011.10.033.

18. McArthur, K., and Kile, B.T. (2018). Apoptotic Caspases: Multiple or Mistaken Identities? Trends Cell Biol 28, 475–493. 10.1016/j.tcb.2018.02.003.

19. Banerjee, U., Girard, J.R., Goins, L.M., and Spratford, C.M. (2019). Drosophila as a Genetic Model for Hematopoiesis. Genetics 211, 367–417. 10.1534/genetics.118.300223.

20. Wood, W., and Martin, P. (2017). Macrophage Functions in Tissue Patterning and Disease: New Insights from the Fly. Dev Cell 40, 221–233. 10.1016/j.devcel.2017.01.001.

21. Kharrat, B., Csordas, G., and Honti, V. (2022). Peeling Back the Layers of Lymph Gland Structure and Regulation. Int J Mol Sci 23. 10.3390/ijms23147767.

22. Mase, A., Augsburger, J., and Bruckner, K. (2021). Macrophages and Their Organ Locations Shape Each Other in Development and Homeostasis - A Drosophila Perspective. Front Cell Dev Biol 9, 630272. 10.3389/fcell.2021.630272.

23. Melcarne, C., Lemaitre, B., and Kurant, E. (2019). Phagocytosis in Drosophila: From molecules and cellular machinery to physiology. Insect Biochem Mol Biol 109, 1–12. 10.1016/j.ibmb.2019.04.002.

24. Coates, J.A., Brooks, E., Brittle, A.L., Armitage, E.L., Zeidler, M.P., and Evans, I.R. (2021). Identification of functionally distinct macrophage subpopulations in Drosophila. Elife 10. 10.7554/eLife.58686.

25. Li, H., Janssens, J., De Waegeneer, M., Kolluru, S.S., Davie, K., Gardeux, V., Saelens, W., David, F.P.A., Brbic, M., Spanier, K., et al. (2022). Fly Cell Atlas: A single-nucleus transcriptomic atlas of the adult fruit fly. Science 375, eabk2432. 10.1126/science.abk2432.

26. Rodrigues, D., Renaud, Y., VijayRaghavan, K., Waltzer, L., and Inamdar, M.S. (2021). Differential activation of JAK-STAT signaling reveals functional compartmentalization in Drosophila blood progenitors. Elife 10. 10.7554/eLife.61409.

27. Mondal, B.C., Shim, J., Evans, C.J., and Banerjee, U. (2014). Pvr expression regulators in equilibrium signal control and maintenance of Drosophila blood progenitors. Elife 3, e03626. 10.7554/eLife.03626.

28. Mondal, B.C., Mukherjee, T., Mandal, L., Evans, C.J., Sinenko, S.A., Martinez-Agosto, J.A., and Banerjee, U. (2011). Interaction between differentiating cell- and niche-derived signals in hematopoietic progenitor maintenance. Cell 147, 1589–1600. 10.1016/j.cell.2011.11.041.

29. Morin-Poulard, I., Vincent, A., and Crozatier, M. (2013). The Drosophila JAK-STAT pathway in blood cell formation and immunity. JAKSTAT 2, e25700. 10.4161/jkst.25700.

30. Shim, J., Mukherjee, T., Mondal, B.C., Liu, T., Young, G.C., Wijewarnasuriya, D.P., and Banerjee, U. (2013). Olfactory control of blood progenitor maintenance. Cell 155, 1141–1153. 10.1016/j.cell.2013.10.032.

31. Shim, J., Mukherjee, T., and Banerjee, U. (2012). Direct sensing of systemic and nutritional signals by haematopoietic progenitors in Drosophila. Nat Cell Biol 14, 394–400. 10.1038/ncb2453.

32. Goyal, M., Tomar, A., Madhwal, S., and Mukherjee, T. (2022). Blood progenitor redox homeostasis through olfaction-derived systemic GABA in hematopoietic growth control in Drosophila. Development 149. 10.1242/dev.199550.

33. Goins, L.M., Girard, J.R., Mondal, B.C., Buran, S., Su, C.C., Tang, R., Biswas, T., and Banerjee, U. (2023). 10.1101/2023.06.29.547151.

34. Owusu-Ansah, E., and Banerjee, U. (2009). Reactive oxygen species prime Drosophila haematopoietic progenitors for differentiation. Nature 461, 537–541. 10.1038/nature08313.

35. Kornepati, A.V.R., Rogers, C.M., Sung, P., and Curiel, T.J. (2023). The complementarity of DDR, nucleic acids and anti-tumour immunity. Nature 619, 475–486. 10.1038/s41586-023-06069-6.

36. Lake, C.M., Holsclaw, J.K., Bellendir, S.P., Sekelsky, J., and Hawley, R.S. (2013). The development of a monoclonal antibody recognizing the Drosophila melanogaster phosphorylated histone H2A variant (gamma-H2AV). G3 (Bethesda) 3, 1539–1543. 10.1534/g3.113.006833.

37. Grigorian, M., DeBruhl, H., and Lipsick, J.S. (2017). The role of variant histone H2AV in Drosophila melanogaster larval hematopoiesis. Development 144, 1441–1449. 10.1242/dev.142729.

38. Evans, C.J., Liu, T., and Banerjee, U. (2014). Drosophila hematopoiesis: Markers and methods for molecular genetic analysis. Methods 68, 242–251. 10.1016/j.ymeth.2014.02.038.

39. Spratford, C.M., Goins, L.M., Chi, F., Girard, J.R., Macias, S.N., Ho, V.W., and Banerjee, U. (2021). Intermediate progenitor cells provide a transition between hematopoietic progenitors and their differentiated descendants. Development 148. 10.1242/dev.200216.

40. Na, H.J., Akan, I., Abramowitz, L.K., and Hanover, J.A. (2020). Nutrient-Driven O-GlcNAcylation Controls DNA Damage Repair Signaling and Stem/Progenitor Cell Homeostasis. Cell Rep 31, 107632. 10.1016/j.celrep.2020.107632.

41. Larsen, B.D., Rampalli, S., Burns, L.E., Brunette, S., Dilworth, F.J., and Megeney, L.A. (2010). Caspase 3/caspase-activated DNase promote cell differentiation by inducing DNA strand breaks. Proc Natl Acad Sci U S A 107, 4230–4235. 10.1073/pnas.0913089107.

42. Gunesdogan, U., Jackle, H., and Herzig, A. (2014). Histone supply regulates S phase timing and cell cycle progression. Elife 3, e02443. 10.7554/eLife.02443.

43. Xu, Y.J., and Leffak, M. (2010). ATRIP from TopBP1 to ATR--in vitro activation of a DNA damage checkpoint. Proc Natl Acad Sci U S A 107, 13561–13562. 10.1073/pnas.1008909107.

44. Blythe, S.A., and Wieschaus, E.F. (2015). Zygotic genome activation triggers the DNA replication checkpoint at the midblastula transition. Cell 160, 1169–1181. 10.1016/j.cell.2015.01.050.

45. Zielke, N., Korzelius, J., van Straaten, M., Bender, K., Schuhknecht, G.F.P., Dutta, D., Xiang, J., and Edgar, B.A. (2014). Fly-FUCCI: A versatile tool for studying cell proliferation in complex tissues. Cell Rep 7, 588–598. 10.1016/j.celrep.2014.03.020.

46. Thacker, S.A., Bonnette, P.C., and Duronio, R.J. (2003). The contribution of E2F-regulated transcription to Drosophila PCNA gene function. Curr Biol 13, 53–58. 10.1016/s0960-9822(02)01400-8.

47. Ding, A.X., Sun, G., Argaw, Y.G., Wong, J.O., Easwaran, S., and Montell, D.J. (2016). CasExpress reveals widespread and diverse patterns of cell survival of caspase-3 activation during development in vivo. Elife 5. 10.7554/eLife.10936.

48. Schott, S., Ambrosini, A., Barbaste, A., Benassayag, C., Gracia, M., Proag, A., Rayer, M., Monier, B., and Suzanne, M. (2017). A fluorescent toolkit for spatiotemporal tracking of apoptotic cells in living Drosophila tissues. Development 144, 3840–3846. 10.1242/dev.149807.

49. Bardet, P.L., Kolahgar, G., Mynett, A., Miguel-Aliaga, I., Briscoe, J., Meier, P., and Vincent, J.P. (2008). A fluorescent reporter of caspase activity for live imaging. Proc Natl Acad Sci U S A 105, 13901–13905. 10.1073/pnas.0806983105.

50. Denton, D., and Kumar, S. (2015). Terminal Deoxynucleotidyl Transferase (TdT)-Mediated dUTP Nick-End Labeling (TUNEL) for Detection of Apoptotic Cells in Drosophila. Cold Spring Harb Protoc 2015, 568–571. 10.1101/pdb.prot086199.

51. Arthurton, L., Nahotko, D.A., Alonso, J., Wendler, F., and Baena-Lopez, L.A. (2020). Non-apoptotic caspase activation preserves Drosophila intestinal progenitor cells in quiescence. EMBO Rep 21, e48892. 10.15252/embr.201948892.

52. Baena-Lopez, L.A., Arthurton, L., Bischoff, M., Vincent, J.P., Alexandre, C., and McGregor, R. (2018). Novel initiator caspase reporters uncover previously unknown features of caspase-activating cells. Development 145. 10.1242/dev.170811.

53. Sun, G., Ding, X.A., Argaw, Y., Guo, X., and Montell, D.J. (2020). Akt1 and dCIZ1 promote cell survival from apoptotic caspase activation during regeneration and oncogenic overgrowth. Nat Commun 11, 5726. 10.1038/s41467-020-19068-2.

54. Csordas, G., Gabor, E., and Honti, V. (2021). There and back again: The mechanisms of differentiation and transdifferentiation in Drosophila blood cells. Dev Biol 469, 135–143. 10.1016/j.ydbio.2020.10.006.

55. Kondo, S., Senoo-Matsuda, N., Hiromi, Y., and Miura, M. (2006). DRONC coordinates cell death and compensatory proliferation. Mol Cell Biol 26, 7258–7268. 10.1128/MCB.00183-06.

56. Muro, I., Berry, D.L., Huh, J.R., Chen, C.H., Huang, H., Yoo, S.J., Guo, M., Baehrecke, E.H., and Hay, B.A. (2006). The Drosophila caspase Ice is important for many apoptotic cell deaths and for spermatid individualization, a nonapoptotic process. Development 133, 3305–3315. 10.1242/dev.02495.

57. Xu, D., Li, Y., Arcaro, M., Lackey, M., and Bergmann, A. (2005). The CARD-carrying caspase Dronc is essential for most, but not all, developmental cell death in Drosophila. Development 132, 2125–2134. 10.1242/dev.01790.

58. Musashe, D.T., Purice, M.D., Speese, S.D., Doherty, J., and Logan, M.A. (2016). Insulin-like Signaling Promotes Glial Phagocytic Clearance of Degenerating Axons through Regulation of Draper. Cell Rep 16, 1838–1850. 10.1016/j.celrep.2016.07.022.

59. Yang, S.A., Portilla, J.M., Mihailovic, S., Huang, Y.C., and Deng, W.M. (2019). Oncogenic Notch Triggers Neoplastic Tumorigenesis in a Transition-Zone-like Tissue Microenvironment. Dev Cell 49, 461–472 e465. 10.1016/j.devcel.2019.03.015.

60. Siegrist, S.E., Haque, N.S., Chen, C.H., Hay, B.A., and Hariharan, I.K. (2010). Inactivation of both Foxo and reaper promotes long-term adult neurogenesis in Drosophila. Curr Biol 20, 643–648. 10.1016/j.cub.2010.01.060.

61. Shinoda, N., Hanawa, N., Chihara, T., Koto, A., and Miura, M. (2019). Dronc-independent basal executioner caspase activity sustains Drosophila imaginal tissue growth. Proc Natl Acad Sci U S A 116, 20539–20544. 10.1073/pnas.1904647116.

62. Evans, C.J., Olson, J.M., Mondal, B.C., Kandimalla, P., Abbasi, A., Abdusamad, M.M., Acosta, O., Ainsworth, J.A., Akram, H.M., Albert, R.B., et al. (2021). A functional genomics screen identifying blood cell development genes in Drosophila by undergraduates participating in a course-based research experience. G3 (Bethesda) 11. 10.1093/g3journal/jkaa028.

63. Purice, M.D., Speese, S.D., and Logan, M.A. (2016). Delayed glial clearance of degenerating axons in aged Drosophila is due to reduced PI3K/Draper activity. Nat Commun 7, 12871. 10.1038/ncomms12871.

64. Kurucz, E., Markus, R., Zsamboki, J., Folkl-Medzihradszky, K., Darula, Z., Vilmos, P., Udvardy, A., Krausz, I., Lukacsovich, T., Gateff, E., et al. (2007). Nimrod, a putative phagocytosis receptor with EGF repeats in Drosophila plasmatocytes. Curr Biol 17, 649–654. 10.1016/j.cub.2007.02.041.

65. Leevers, S.J., Weinkove, D., MacDougall, L.K., Hafen, E., and Waterfield, M.D. (1996). The Drosophila phosphoinositide 3-kinase Dp110 promotes cell growth. EMBO J 15, 6584–6594.

66. Santabarbara-Ruiz, P., Lopez-Santillan, M., Martinez-Rodriguez, I., Binagui-Casas, A., Perez, L., Milan, M., Corominas, M., and Serras, F. (2015). ROS-Induced JNK and p38 Signaling Is Required for Unpaired Cytokine Activation during Drosophila Regeneration. PLoS Genet 11, e1005595. 10.1371/journal.pgen.1005595.

67. Toshniwal, A.G., Gupta, S., Mandal, L., and Mandal, S. (2019). ROS Inhibits Cell Growth by Regulating 4EBP and S6K, Independent of TOR, during Development. Dev Cell 49, 473–489 e479. 10.1016/j.devcel.2019.04.008.

68. Santabarbara-Ruiz, P., Esteban-Collado, J., Perez, L., Viola, G., Abril, J.F., Milan, M., Corominas, M., and Serras, F. (2019). Ask1 and Akt act synergistically to promote ROS-dependent regeneration in Drosophila. PLoS Genet 15, e1007926. 10.1371/journal.pgen.1007926.

69. Uhlirova, M., and Bohmann, D. (2006). JNK- and Fos-regulated Mmp1 expression cooperates with Ras to induce invasive tumors in Drosophila. EMBO J 25, 5294–5304. 10.1038/sj.emboj.7601401.

70. Girard, J.R., Goins, L.M., Vuu, D.M., Sharpley, M.S., Spratford, C.M., Mantri, S.R., and Banerjee, U. (2021). Paths and pathways that generate cell-type heterogeneity and developmental progression in hematopoiesis. Elife 10. 10.7554/eLife.67516.

71. Sykiotis, G.P., and Bohmann, D. (2008). Keap1/Nrf2 signaling regulates oxidative stress tolerance and lifespan in Drosophila. Dev Cell 14, 76–85. 10.1016/j.devcel.2007.12.002.

72. Larsen, B.D., and Sorensen, C.S. (2017). The caspase-activated DNase: apoptosis and beyond. FEBS J 284, 1160–1170. 10.1111/febs.13970.

73. Enari, M., Sakahira, H., Yokoyama, H., Okawa, K., Iwamatsu, A., and Nagata, S. (1998). A caspase-activated DNase that degrades DNA during apoptosis, and its inhibitor ICAD. Nature 391, 43–50. 10.1038/34112.

74. Ha, H.J., and Park, H.H. (2022). Molecular basis of apoptotic DNA fragmentation by DFF40. Cell Death Dis 13, 198. 10.1038/s41419-022-04662-7.

75. Choi, J.Y., Qiao, Q., Hong, S.H., Kim, C.M., Jeong, J.H., Kim, Y.G., Jung, Y.K., Wu, H., and Park, H.H. (2017). CIDE domains form functionally important higher-order assemblies for DNA fragmentation. Proc Natl Acad Sci U S A 114, 7361–7366. 10.1073/pnas.1705949114.

76. Mukae, N., Yokoyama, H., Yokokura, T., Sakoyama, Y., Sakahira, H., and Nagata, S. (2000). Identification and developmental expression of inhibitor of caspase-activated DNase (ICAD) in Drosophila melanogaster. J Biol Chem 275, 21402–21408. 10.1074/jbc.M909611199.

77. Lee, P.T., Zirin, J., Kanca, O., Lin, W.W., Schulze, K.L., Li-Kroeger, D., Tao, R., Devereaux, C., Hu, Y., Chung, V., et al. (2018). A gene-specific T2A-GAL4 library for Drosophila. Elife 7. 10.7554/eLife.35574.

78. Tarayrah-Ibraheim, L., Maurice, E.C., Hadary, G., Ben-Hur, S., Kolpakova, A., Braun, T., Peleg, Y., Yacobi-Sharon, K., and Arama, E. (2021). DNase II mediates a parthanatos-like developmental cell death pathway in Drosophila primordial germ cells. Nat Commun 12, 2285. 10.1038/s41467-021-22622-1.

79. Yacobi-Sharon, K., Namdar, Y., and Arama, E. (2013). Alternative germ cell death pathway in Drosophila involves HtrA2/Omi, lysosomes, and a caspase-9 counterpart. Dev Cell 25, 29–42. 10.1016/j.devcel.2013.02.002.

80. Shravage, B.V., Hill, J.H., Powers, C.M., Wu, L., and Baehrecke, E.H. (2013). Atg6 is required for multiple vesicle trafficking pathways and hematopoiesis in Drosophila. Development 140, 1321–1329. 10.1242/dev.089490.

81. Kadandale, P., Stender, J.D., Glass, C.K., and Kiger, A.A. (2010). Conserved role for autophagy in Rho1-mediated cortical remodeling and blood cell recruitment. Proc Natl Acad Sci U S A 107, 10502–10507. 10.1073/pnas.0914168107.

82. Zhang, Y., Morgan, M.J., Chen, K., Choksi, S., and Liu, Z.G. (2012). Induction of autophagy is essential for monocyte-macrophage differentiation. Blood 119, 2895–2905. 10.1182/blood-2011-08-372383.

83. Jacquel, A., Obba, S., Boyer, L., Dufies, M., Robert, G., Gounon, P., Lemichez, E., Luciano, F., Solary, E., and Auberger, P. (2012). Autophagy is required for CSF-1-induced macrophagic differentiation and acquisition of phagocytic functions. Blood 119, 4527–4531. 10.1182/blood-2011-11-392167.

84. Nezis, I.P., Shravage, B.V., Sagona, A.P., Lamark, T., Bjorkoy, G., Johansen, T., Rusten, T.E., Brech, A., Baehrecke, E.H., and Stenmark, H. (2010). Autophagic degradation of dBruce controls DNA fragmentation in nurse cells during late Drosophila melanogaster oogenesis. J Cell Biol 190, 523–531. 10.1083/jcb.201002035.

85. Jipa, A., Vedelek, V., Merenyi, Z., Urmosi, A., Takats, S., Kovacs, A.L., Horvath, G.V., Sinka, R., and Juhasz, G. (2021). Analysis of Drosophila Atg8 proteins reveals multiple lipidation-independent roles. Autophagy 17, 2565–2575. 10.1080/15548627.2020.1856494.

86. Rahman, A., Lorincz, P., Gohel, R., Nagy, A., Csordas, G., Zhang, Y., Juhasz, G., and Nezis, I.P. (2022). GMAP is an Atg8a-interacting protein that regulates Golgi turnover in Drosophila. Cell Rep 39, 110903. 10.1016/j.celrep.2022.110903.

87. Klionsky, D.J., Abdel-Aziz, A.K., Abdelfatah, S., Abdellatif, M., Abdoli, A., Abel, S., Abeliovich, H., Abildgaard, M.H., Abudu, Y.P., Acevedo-Arozena, A., et al. (2021). Guidelines for the use and interpretation of assays for monitoring autophagy (4th edition)(1). Autophagy 17, 1–382. 10.1080/15548627.2020.1797280.

88. Tattikota, S.G., Cho, B., Liu, Y., Hu, Y., Barrera, V., Steinbaugh, M.J., Yoon, S.H., Comjean, A., Li, F., Dervis, F., et al. (2020). A single-cell survey of Drosophila blood. Elife 9. 10.7554/eLife.54818.

89. Gyoergy, A., Roblek, M., Ratheesh, A., Valoskova, K., Belyaeva, V., Wachner, S., Matsubayashi, Y., Sanchez-Sanchez, B.J., Stramer, B., and Siekhaus, D.E. (2018). Tools Allowing Independent Visualization and Genetic Manipulation of Drosophila melanogaster Macrophages and Surrounding Tissues. G3 (Bethesda) 8, 845–857. 10.1534/g3.117.300452.

90. McPhee, C.K., Logan, M.A., Freeman, M.R., and Baehrecke, E.H. (2010). Activation of autophagy during cell death requires the engulfment receptor Draper. Nature 465, 1093–1096. 10.1038/nature09127.

91. Nagarkar-Jaiswal, S., DeLuca, S.Z., Lee, P.T., Lin, W.W., Pan, H., Zuo, Z., Lv, J., Spradling, A.C., and Bellen, H.J. (2015). A genetic toolkit for tagging intronic MiMIC containing genes. Elife 4. 10.7554/eLife.08469.

92. Melcarne, C., Ramond, E., Dudzic, J., Bretscher, A.J., Kurucz, E., Ando, I., and Lemaitre, B. (2019). Two Nimrod receptors, NimC1 and Eater, synergistically contribute to bacterial phagocytosis in Drosophila melanogaster. FEBS J 286, 2670–2691. 10.1111/febs.14857.

93. Amcheslavsky, A., Lindblad, J.L., and Bergmann, A. (2020). Transiently “Undead” Enterocytes Mediate Homeostatic Tissue Turnover in the Adult Drosophila Midgut. Cell Rep 33, 108408. 10.1016/j.celrep.2020.108408.

94. Fujisawa, Y., Shinoda, N., Chihara, T., and Miura, M. (2020). ROS Regulate Caspase-Dependent Cell Delamination without Apoptosis in the Drosophila Pupal Notum. iScience 23, 101413. 10.1016/j.isci.2020.101413.

95. White, K., Arama, E., and Hardwick, J.M. (2017). Controlling caspase activity in life and death. PLoS Genet 13, e1006545. 10.1371/journal.pgen.1006545.

96. Jacquel, A., Benikhlef, N., Paggetti, J., Lalaoui, N., Guery, L., Dufour, E.K., Ciudad, M., Racoeur, C., Micheau, O., Delva, L., et al. (2009). Colony-stimulating factor-1-induced oscillations in phosphatidylinositol-3 kinase/AKT are required for caspase activation in monocytes undergoing differentiation into macrophages. Blood 114, 3633–3641. 10.1182/blood-2009-03-208843.

97. Weavers, H., Evans, I.R., Martin, P., and Wood, W. (2016). Corpse Engulfment Generates a Molecular Memory that Primes the Macrophage Inflammatory Response. Cell 165, 1658–1671. 10.1016/j.cell.2016.04.049.

98. Machour, F.E., and Ayoub, N. (2020). Transcriptional Regulation at DSBs: Mechanisms and Consequences. Trends Genet 36, 981–997. 10.1016/j.tig.2020.01.001.

99. Cho, B., Yoon, S.H., Lee, D., Koranteng, F., Tattikota, S.G., Cha, N., Shin, M., Do, H., Hu, Y., Oh, S.Y., et al. (2020). Single-cell transcriptome maps of myeloid blood cell lineages in Drosophila. Nat Commun 11, 4483. 10.1038/s41467-020-18135-y.

100. Nano, M., Mondo, J.A., Harwood, J., Balasanyan, V., and Montell, D.J. (2023). Cell survival following direct executioner-caspase activation. Proc Natl Acad Sci U S A 120, e2216531120. 10.1073/pnas.2216531120.

101. Yu, J.S., and Cui, W. (2016). Proliferation, survival and metabolism: the role of PI3K/AKT/mTOR signalling in pluripotency and cell fate determination. Development 143, 3050–3060. 10.1242/dev.137075.

102. Yang, J.Y., Michod, D., Walicki, J., Murphy, B.M., Kasibhatla, S., Martin, S.J., and Widmann, C. (2004). Partial cleavage of RasGAP by caspases is required for cell survival in mild stress conditions. Mol Cell Biol 24, 10425–10436. 10.1128/MCB.24.23.10425-10436.2004.

103. Shemorry, A., Harnoss, J.M., Guttman, O., Marsters, S.A., Komuves, L.G., Lawrence, D.A., and Ashkenazi, A. (2019). Caspase-mediated cleavage of IRE1 controls apoptotic cell commitment during endoplasmic reticulum stress. Elife 8. 10.7554/eLife.47084.

104. Cho, U.H., and Hetzer, M.W. (2023). Caspase-mediated nuclear pore complex trimming in cell differentiation and endoplasmic reticulum stress. Elife 12. 10.7554/eLife.89066.

105. Al-Khalaf, M.H., Blake, L.E., Larsen, B.D., Bell, R.A., Brunette, S., Parks, R.J., Rudnicki, M.A., McKinnon, P.J., Jeffrey Dilworth, F., and Megeney, L.A. (2016). Temporal activation of XRCC1-mediated DNA repair is essential for muscle differentiation. Cell Discov 2, 15041. 10.1038/celldisc.2015.41.

106. Benada, J., Alsowaida, D., Megeney, L.A., and Sorensen, C.S. (2023). Self-inflicted DNA breaks in cell differentiation and cancer. Trends Cell Biol 33, 850–859. 10.1016/j.tcb.2023.03.002.

107. Spencer, S.L., and Sorger, P.K. (2011). Measuring and modeling apoptosis in single cells. Cell 144, 926–939. 10.1016/j.cell.2011.03.002.

108. Larsen, B.D., Benada, J., Yung, P.Y.K., Bell, R.A.V., Pappas, G., Urban, V., Ahlskog, J.K., Kuo, T.T., Janscak, P., Megeney, L.A., et al. (2022). Cancer cells use self-inflicted DNA breaks to evade growth limits imposed by genotoxic stress. Science 376, 476–483. 10.1126/science.abi6378.

109. Dehingia, B., Milewska, M., Janowski, M., and Pekowska, A. (2022). CTCF shapes chromatin structure and gene expression in health and disease. EMBO Rep 23, e55146. 10.15252/embr.202255146.

110. Puc, J., Aggarwal, A.K., and Rosenfeld, M.G. (2017). Physiological functions of programmed DNA breaks in signal-induced transcription. Nat Rev Mol Cell Biol 18, 471–476. 10.1038/nrm.2017.43.

111. Simmons, J.R., Kemp, J.D.J., and Labrador, M. (2023). 10.1101/2023.05.31.543180.

112. Bekkering, S., Dominguez-Andres, J., Joosten, L.A.B., Riksen, N.P., and Netea, M.G. (2021). Trained Immunity: Reprogramming Innate Immunity in Health and Disease. Annu Rev Immunol 39, 667–693. 10.1146/annurev-immunol-102119-073855.

